# Clustering of death receptor for apoptosis using nanoscale patterns of peptides

**DOI:** 10.1101/2020.11.16.359729

**Authors:** Yang Wang, Igor Baars, Ferenc Fördös, Björn Högberg

## Abstract

The nanoscale spatial organization of transmembrane tumor necrosis factor (TNF) receptors has been implied as a regulator of cellular fate. Accordingly, molecular tools that can induce specific arrangements of these receptors on cell surfaces would give us an opportunity to study these effects in detail. To achieve this, we introduce DNA origami nanostructures, that precisely scaffold the patterning of TNF-related apoptosis-inducing ligand (TRAIL)-mimicking peptides at nanoscale level. Stimulating human breast cancer cells with these patterns, we find that around 5 nm is the critical inter-ligand distance of hexagonally patterned peptides to induce death receptor clustering and a resulting apoptosis. We thus offer a strategy to reverse the non-efficacy of current ligand- and antibody-based methods for TNF superfamily (TNFRSF) activation.

## Introduction

Because the tumor necrosis factor (TNF) receptor superfamily (TNFRSF) plays important roles in cell proliferation, cell death, immune regulation and morphogenesis, it has been extensively targeted for disease treatment^1–5^. Although structural information of TNFRSF, corresponding ligands and even the receptor-ligand complexes have been quite thoroughly characterized^6, 7^, molecular tools and drugs that can effectively trigger TNFRSF signaling are still missing^3, 5^. Currently available agonists usually fail to work as expected^8^. A typical example is the TNF-related apoptosis inducing ligand (TRAIL). TRAIL can recognize and bind to death receptors (DR), including death receptor 4 (DR4) and death receptor 5 (DR5)^9^. Human TRAIL (Dulanermin) and DR4/5 agonistic monoclonal antibodies (e.g., Mapatumumab and Lexatumumab) have been used as anti-cancer therapeutics since the mid-1990s^10, 11^. Randomized control trials have however recently shown that their efficiency in terms of survival benefits are lacking^12, 13^. Studies of TRAIL-DR5 complexes on cells displaying apoptosis have revealed that the transmembrane domain of DR5 formed higher-order structures through clustering^14^. Thus, one potential reason for the lack of efficacy could be that death receptor triggering via the natural ligand/antibody-receptor binding mechanism, might not be strong enough to cause receptor clustering. To explore this, anti-TRAIL antibodies, which can crosslink TRAIL, were used together with TRAIL to promote the formation of DR5 clusters^15, 16^. This strategy improved the apoptosis of cancer cells. Another strategy was to covalently multimerize TRAIL or TRAIL-mimicking peptide on peptide-, dextran- or graphene-based scaffolds^17–20^, which was also demonstrated to be efficacious. However, using these strategies it is typically not possible to precisely control the nanoscale spatial presentation of the proteins or peptides. By conjugating ligands onto surfaces with pre-patterned nanodot arrays, other groups have achieved a nanoscale arrangement of TNF with spacings between 58 and 290 nm^21^. Cell culture on those surfaces showed a dependence on inter-ligand distances, revealing the importance of inter-ligand distance control for efficient death receptor activation. Nevertheless, the achievable smallest inter-dot distance was the size of nanodots themselves, making for example sub-10 nm inter-dot arrangement a challenge. Also, note that in display experiments where a surface is covered in ligands, both the number and spatial separation between them are varied at the same time. Consequently, surfaces with differently spaced ligands would also display different overall ligand amounts at the surface-cell interface, which could also affect cell activity, giving potentially confounded interpretations. On top of this, clinical translations of surface patterning methods are typically limited as the path from patterning large planar surfaces to patterning biocompatible nanoparticles is not straightforward.

In contrast, DNA origami nanostructures^22–29^ offer both a programmable way to precisely display biomolecule nanopatterns from monodisperse particles, and through this a way to display these to cells from the solution phase. This allows us to vary the separation of ligands independently from the total dosage, or total concentration of ligands and further allows us to focus solely on the spatial separation between ligands on the nanoscale. Thanks to its spatial addressability^30–33^, varying nanopatterns of ephrin-A5^34, 35^, caspase-9 variant^36^, antigens of human IgGs and IgMs^37^, immunogen eOD-GT8^38^ and Fas ligands^39^ on DNA origami nanostructures have been studied, showing an increasing importance for biomedical applications.

## Results and discussions

To investigate the effect of differing ligand pattern sizes on death receptors, we prepared two versions of DNA origami as templates: a single-layer wireframe (W) flat sheet (Fig.1a) and a double-layer square lattice (L) style flat sheet (Fig.1b; Extended Fig.1, 2). Sharp electrophoresis bands of structures (before and after purification) on 2% agarose gels (Extended Fig.3) and expected structural characteristics under atomic force microscopy (AFM) (Fig.1c, d) indicate successful structure preparations. We produced a collection of structures displaying protruding 5’ single-stranded DNA (ssDNA) handles for subsequent hybridization of ssDNA-ligand conjugates.

**Fig.1.**
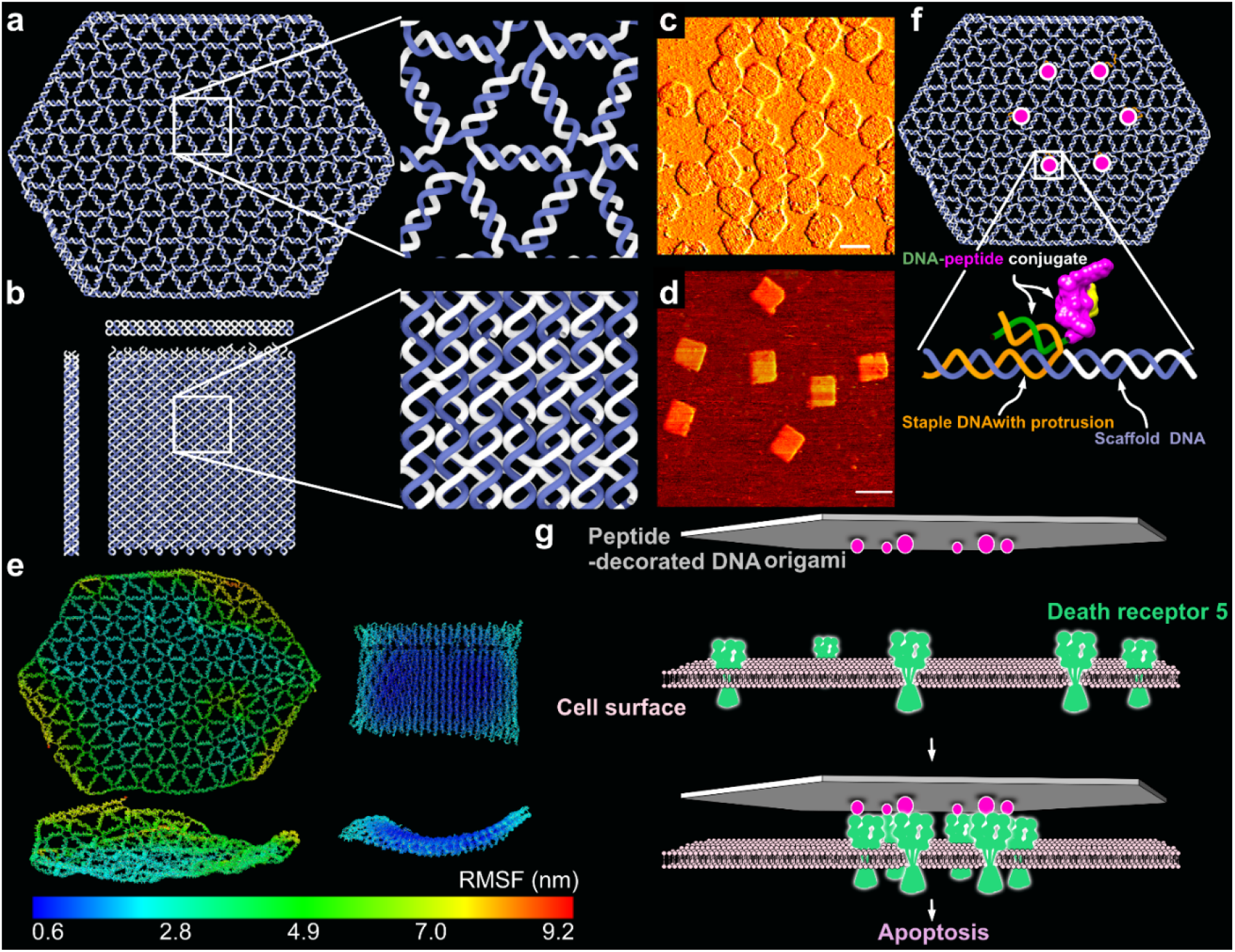
Structure design and characterization. **a**, Stylized renderings of the single-layer W-structure (top view), together with highlight of its DNA arrangements details. **b**, Stylized renderings of the double-layer L-structure (side view, front view and top view), together with highlight of its DNA arrangements details. The scaffold DNA is colored in dark blue, and staples are in gray. **c** and **d**, The two types of structures imaged by atomic force microscopy (AFM). Scale bars are 100 nm. **e**, Computed mean structures and RMSF of the W- and L-structures using oxDNA, top and front views. **f**, Schematic illustration of how the TRAIL-mimicking peptides are attached to the DNA origamis (same scheme in both W and L). **g**, Schematic illustration of how breast cancer cell apoptosis can be triggered by peptide patterns on DNA origami templates by inducing clustering of death receptor 5.

Because of the different structural design methods for W and L, they are expected to possess differences in local rigidity and propensity for thermal fluctuation. The mean structures, which were computed from oxDNA molecular dynamics simulation^40, 41^ in 500 mM Na^+^ (simulation parameter), showed that the L-type structure tended to stay flatter than the W types(Fig.1e). Root mean square fluctuation (RMSF) values of the Ws are estimated to be around 5-time higher than those of the Ls (Fig.1e), further validating its higher local structural flexibility. These differences are a consequence of the design schemes that can be attributed to the fact that DNA helices in L (Fig.1b) are more restrained through multiple connections to other helices in the structures as is common in latticebased 3D origami. This analysis and similar previous investigations on wireframe structures^42^ suggest that the displayed separations on the W-structures are probably fluctuating more during experiments than the corresponding ones on the L-structures.

Homo-trimeric TRAIL is the natural ligand of pre-ligand DR5 association, although it still not clear if the oligomeric state of the pre-ligand DR5 association is trimeric (Extended Fig.7a)^43^ or dimeric (Extended Fig.7e)^7,14,44^. We used a small cyclic peptide composed of 17 amino acids (Extended Fig.5). This peptide has shown its ability to compete the binding of homo-trimeric TRAIL to pre-ligand DR5 association and thus mimic TRAIL’s functions^45,46^. The rationale for this was two-fold: 1) The much smaller spatial dimension of the peptide, which has its Van der Waals radius at ~1.5 nm (Extended Fig.4a), could allow sub-10nm interligand spacing, while the spatial dimension of homo-trimeric TRAIL (PDB ID: 1DG6) itself is already around 10 nm (Extended Fig.4b). 2) Ease of DNA conjugation: While conjugating a protein like TRAIL monomer/trimer with ssDNA could potentially result in multiple modifications per protein or complex. This would in turn risk having one protein occupy more than one binding site of DNA origami, impeding the formation of expected protein patterns. In contrast, we could easily achieve one ssDNA per peptide conjugation by chemically targeting one pre-functionalized site of the peptide.

We produced the ssDNA-peptide conjugate via click chemistry between an azide functionalized on lysine located at the C-terminal of the peptide and a dibenzocyclooctyne (DBCO) modification at the 5-prime end of the ssDNA (Extended Fig.5). On native polyacrylamide gels, the conjugate had a slower electrophoretic migration than the peptide itself, which were clearly visualized by fluorescent DNA labelling and peptide staining (Extended Fig.6), verifying the reliability of this conjugation method. Importantly, we used a strategy where the conjugation site is on the 5-prime end and binding to a 5-prime end protrusion, thus constraining the peptide close to the site on the DNA origami where the protrusion originates (Fig.1f). Proceeding this way, we avoid a large distance being introduced by the hybridization.

Previous studies on DR5 clusters on apoptotic cells has led to an hypothesis that nanoscale hexagonal DR5 network formed through the dimerization of DR5 trimers (Extended Fig.7b, c) can directly result in apoptosis^47^. More recent studies have then shown that the signaling driver is more likely due to the formation of higher-order transmembrane helix (TMH) structures via the trimerization of DR5 dimers (Extended Fig.7f, g). Similarly to earlier hypothesis though, the potential clustering network mediated by DR5 TMH would again be presented by a hexagonal pattern^14^. Pre-ligand binding, however, this formation was inhibited by the extracellular domain of DR5. Antibody AMG655 can promote the homotrimeric TRAIL’s efficacies on DR5 clustering and antitumor activity. Position modeling of the crystallographically-decoded TRAIL-DR5-AMG 655 Fab ternary complex further emphasized the importance of the hexagonal honeycomb DR5 pattern on apoptosis (Extended Fig.7d, h)^15^. Following this, we aimed to develop a method to precisely induce this active hexagonally organized DR5 network in breast cancer cells, irrespective of whether the actual molecular mechanism to trigger apoptosis is dimerization of DR5 trimers or trimerization of DR5 dimers (Fig.1g).

We thus decided to investigate patterns of peptides displayed in hexagons, where all pattern had sizes less than 50 nm in intra-peptide spacing. We investigated 5.7-nm (W6), 9.43-nm (W9), 15.8-nm (W16), 18.8-nm (W19) and 25.5-nm (W26) peptide patterns on the wireframe (W) structures (Extended Fig.8), and 6.3-nm (L6), 11.1-nm (L11) peptide patterns on the square lattice (L) structures (Extended Fig.9, 10). The nominal distances used in the naming of the structures are taken from the mean distance of the 6 individual nucleotide-to-nucleotide distances (corresponding to the 6 edges of the hexagons) on the DNA origami from oxDNA molecular dynamics simulations (Extended Fig.11). First, electrophoresis of these structures on agarose gel show clean monomeric products (Extended Fig.12, 13). Peptide attachment does not appear to affect the gel mobility, which could be expected as the molecular weight increase (from 5.02 MDa to 5.06 MDa), resulting from hybridizing the peptide-DNA conjugates to the origamis, is minute (0.80% increase). Instead, to verify the correct localization of peptides on structures, we performed a DNA-PAINT experiment: For this analysis only, we inserted 9 extra nucleotides (NT) between the peptide and the origami-hybridizing region close to the 5’ of the DNA in the ssDNA-peptide conjugates (Fig. 2a). These 9 NT regions were used as docking sites for transient binding of Cy3-labeled DNA imager strands. This transient binding process was then imaged using DNA-PAINT.^48–50^ Imaging results showed the expected hexagonal patterns (Fig. 2b, Extended Fig.14-23), indicating an overall correct localization of the peptides on DNA origami. Not all structures in each sample showed a 6-spot hexagonal pattern, and the distribution analysis showed individual site occupancy rates of 49% to 76% depending on structure (Extended Fig.24). The occupancy rate was however estimated to be higher following gel analysis (see below) and it could be that the analysis is underestimating the incorporation possibly due to the 9nt PAINT-imaging sites getting sterically blocked from PAINT-probe binding as they lie sandwiched between the protruding site dsDNA and the peptide itself. The pattern size distributions calculated from DNA-PAINT data appear smaller, in particular for the larger patterns, than the sizes in design and oxDNA simulations (Extended Fig.25). This is to some degree expected as the wireframe structures have a higher sensitivity for global deformations (which would primarily impact the larger patterns) as the structures land on the biotinylated surface during PAINT imaging. Note that this strategy is not imaging empty sites, but importantly we are estimating the actual incorporation of DNA-peptide conjugates by targeting the conjugate strand itself with the PAINT probes (Fig. 2a).

**Fig.2.**
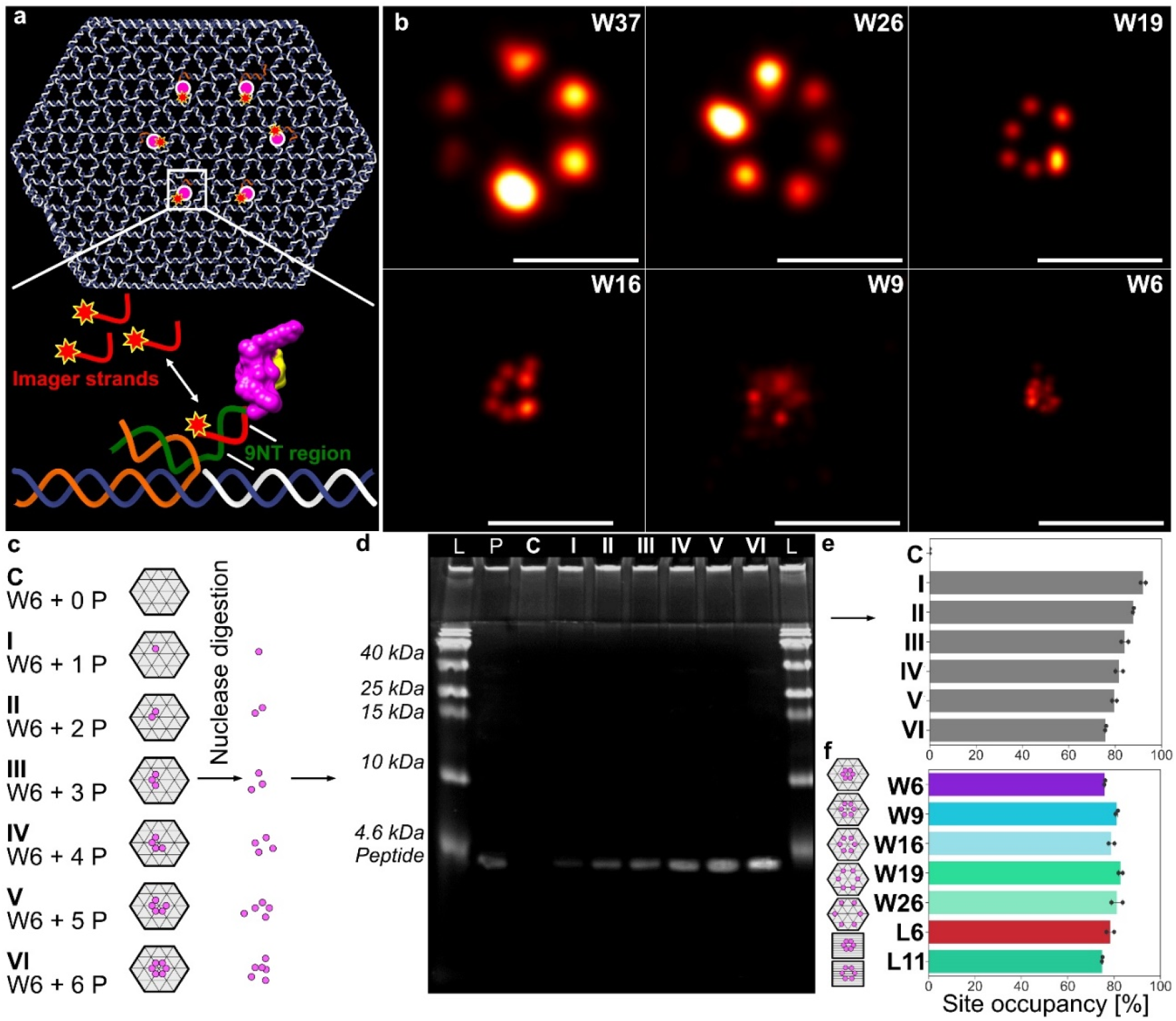
Peptide pattern imaging and quantification. **a**, Schematic illustration of how Cy3-labeled imager strands transiently bind to the 9 extra nucleotides (NT) region between the peptide and the origami-hybridizing region of the DNA in the ssDNA-peptide conjugate in DNA PAINT experiment. **b**, representative DNA-PAINT images of the differently-sized peptide patterns on DNA origamis. Scale bars are 50 nm. W37 is a 37-nm peptide pattern on a wireframe (W) structure used exclusively as a reference during PAINT imaging and analysis. **c** and **d**, W5 structures decorated with varying numbers of peptides were treated with DNase I to completely digest the DNA origami template. The samples were then run on an SDS-PAGE gel, and imaged via Silver staining. L = Protein ladder, P = Peptide alone, C = Empty origami control, I-VI = Decorated origamis according to **c**. **e**, Dosiometry of the gel images plotted as estimated protrusion site occupancy by ssDNA-peptide conjugate on W6 (with varying number of protruding ssDNA sites). **f,** Similar analysis of all estimated calculated protrusion site occupancy by ssDNA-peptide conjugate on the different DNA origamis (with 6 protruding DNA sites) used in the study.

To further verify the stoichiometry of peptides on each structure, we performed a protein gel analysis after a DNAse treatment. Briefly, fully assembled and purified test structures including the positioned peptides, were first incubated with DNAse I (Fig. 2c; Extended Fig.26a). Following this, we ran the resulting degradation products on SDS-PAGE gels and stained for peptides. After which we performed dosimetry analysis on the lanes to estimate the total peptide content in the samples. The analysis showed a clear correlation between the number of protruding DNA sites per structure and the silver staining intensity of peptide bands (Fig. 2d; Extended Fig.26b), corroborating a close match between the designed stoichiometry and the experimental implementation of the structures. On both W and L structures, we estimate that the protruding DNA site occupancy by ssDNA-peptide conjugate was decreasing, from around 93% (with 1 protrusion site) to around 75% (with 6 protrusion sites), with the increasing of the number of protrusion site (Fig. 2e; Extended Fig.26c). This could probably be attributed to a combination of charge- and steric-hindrance effects. In the target hexagonal peptide patterns, the estimated percentage of site occupancy is between 75% and 80% (Fig. 2f), which is slightly higher than the corresponding values from the DNA PAINT assay (Extended Fig.24).

Human cancer cells can respond to TRAIL-treatment very differently^51^. When it comes to breast cancer cells, the triple-negative mesenchymal ones, to which MDA-MB-231 cells belong, were found to be sensitive to TRAIL.^52^ However, those having receptors for estrogen, which include MCF-7 cells, appear to be largely resistant to TRAIL. Those with Her-2 upregulation, including the SK-BR-3 cells, also appear to have a low sensitivity to TRAIL treatment. Although the expression level of death receptors could partially be linked to this phenomenon^53^, and intracellular negative feedbacks on TRAIL response were revealed in certain cells^54^, the actual mechanisms behind this discrepancy are still not well understood. We used MDA-MB-231, MCF-7 and SK-BR-3 cell lines to cover all these three types of human breast cancer.

To visualize DR5 clusters, we firstly established GFP-DR5-expressing cells by plasmid transfection (Fig.3a). We then treated these cells with DNA origami presenting TRAIL-mimicking peptide patterns, following by imaging cells under confocal microscopy. On all cell lines, it showed that (Fig.3b-e): neither did peptide itself nor did DNA origami structures themselves have observable effects on DR5 clustering; for peptide patterns, W6 and W9 successfully caused DR5 clusters, while W16, W19 or W26 didn’t. This size-dependent effect indicated that, to trigger the process of DR5 clustering, peptides need to be patterned closely enough, and around 10 nm seemed to be the critical distance. We also observed significant differences between cells treated with W6 and cells treated with W9.We then further investigated this with the stiffer structure—L6 and L11. The results showed that (Fig.3b-e), as being more effective than W6, L6 successfully triggered DR5 clustering. Interestingly, being different from W9, L11 failed to cause DR5 clustering. This could be explained by the higher structural flexibility of W. As the result, spatial positions of peptides can highly fluctuate within the wireframe structure, eliminating the subtle pattern size difference to a certain extent.

**Fig.3.**
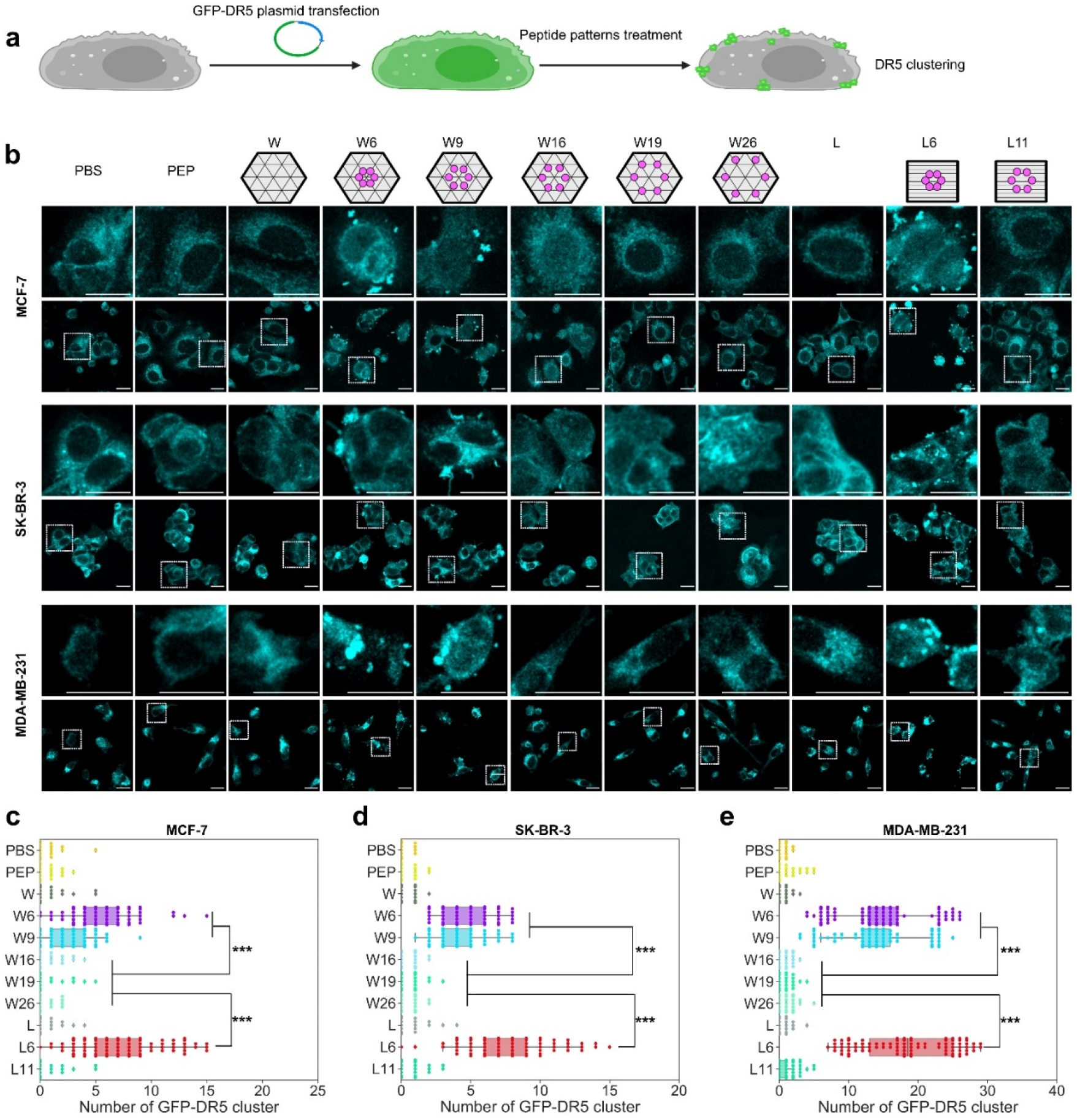
Peptide patterns-induced GFP-DR5 clusters. **a,** Experimental workflow. **b**, GFP-DR5 (Cyan) clusters in breast cancer cells with 4 hours’ treatments of 2-nM DNA origami structures or 12-nM peptides. Scale bars are 20 μm. The enlarged figures above correspond to the indicated areas in figures below. **c-e**, GFP-DR5 cluster counting of MCF-7 cells (**c**), SK-BR-3 cells (**d**) and MDA-MB-231 cells (**e**). Each point stands for the number of GFP-DR5 cluster for one cell (n = 100 cells). PBS stands for phosphate-buffered saline, and PEP stands for peptide. p* < 0.05. p** < 0.01.

In both W and L cases, notably, for all three cell lines, treated cells having GFP-DR5 clusters showed high colocalizations with Cy5-labled DNA origami (Fig.4a-d). We also found that, on MCF-7 cells, the extent of DR5 clustering decreased with respect to the deduction of peptide per L6 (Extended Fig.27), validating the importance of the pattern in hexagon. With these results together, we concluded that DR5 clustering is a precisely controlled cellular process, whose effective triggering needs around 5 nm-spaced hexagonal ligand patterns.

**Fig.4.**
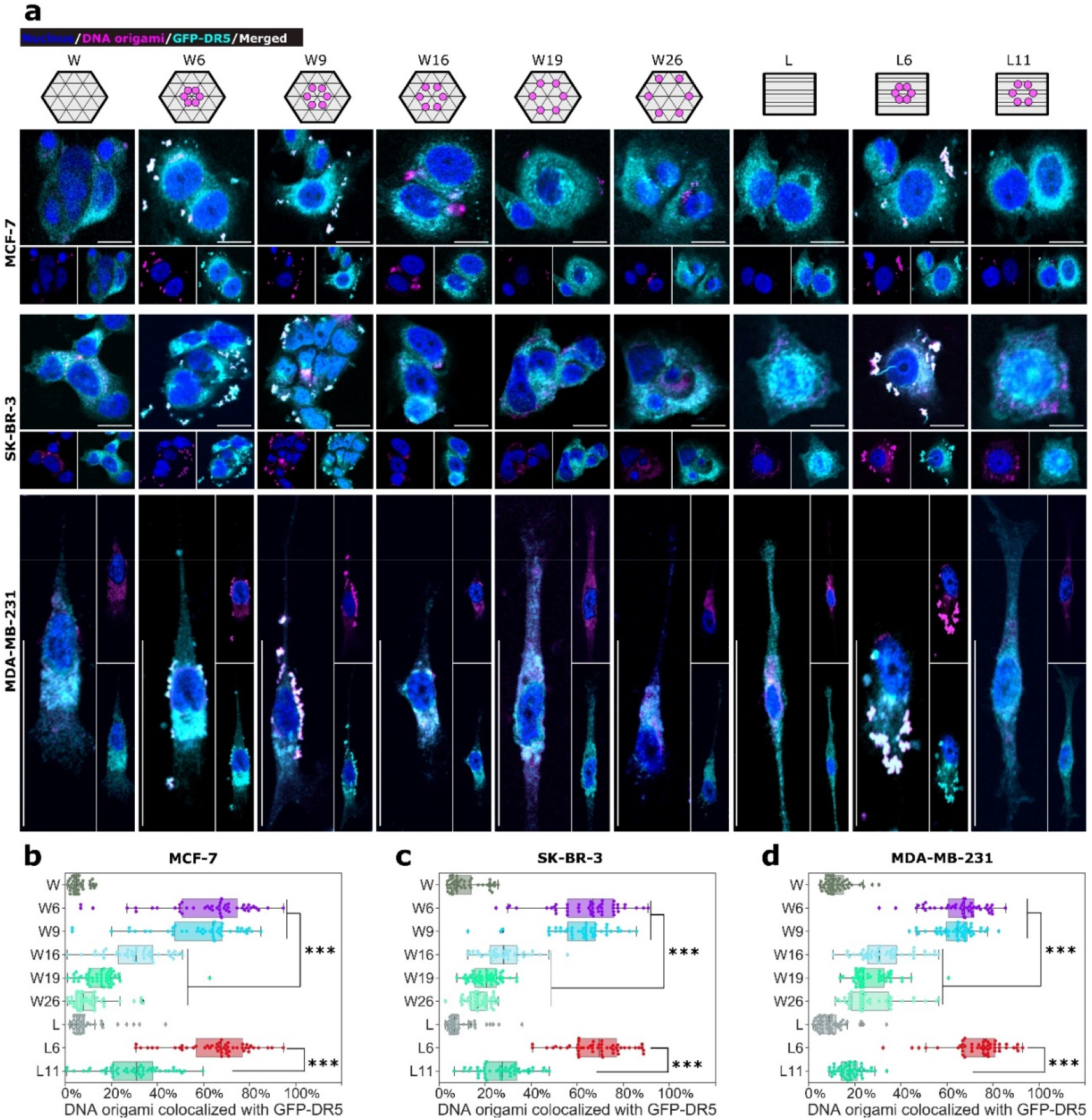
DNA origamis and GFP-DR5 clusters co-localization. **a**, Localization of DNA origami (Magenta) relative to GFP-DR5 (Cyan) clusters. The cells were treated with 2-nM DNA origami structures or 12-nM peptides for 4 hours. Scale bars are 20 μm. **b-d**, Percentage of DNA origami colocalized with GFP-DR5 clusters (n = 50 fields.) from MCF-7 (**b**), SK-BR-3 (**c**) and MDA-MB-231 (**d**) cells. p*** < 0.001.

Based on these DR5 clustering results, we then measured the apoptosis in nontransfected cell lines. On all three cell lines (Fig.5), W6 or W9 induced significantly more apoptotic and apoptosis-resulted necrotic cells than W16, W19 and W26. For cells treated with L6 or L11, however, only L6 showed a strong apoptosis-inducing capability. Empty DNA origami or peptide itself showed no effect.

**Fig.5.**
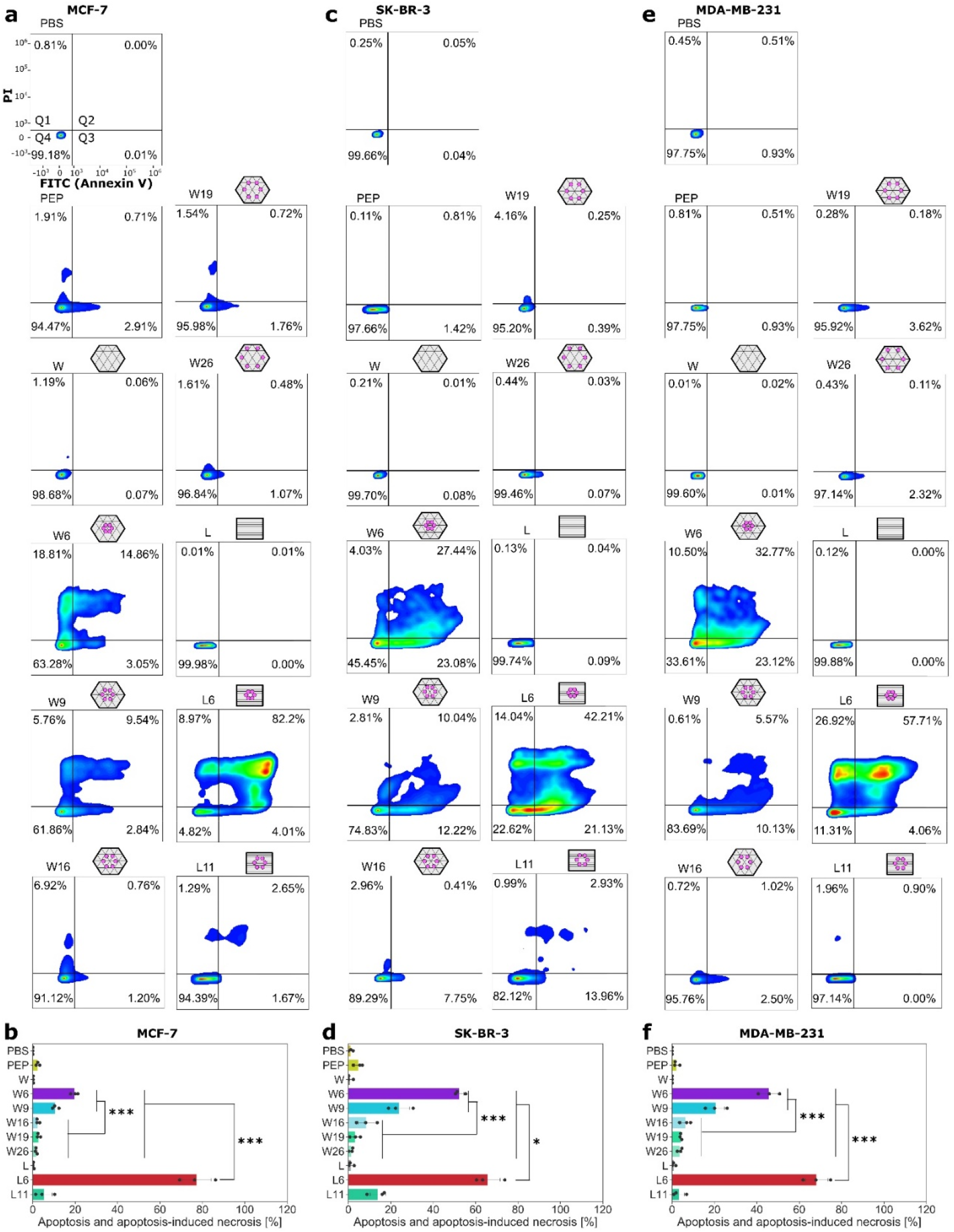
Apoptosis of cells treated with peptide patterns. **a**, **c**, **e**, Representative MCF-7 (**a**), SK-BR-3 (**c**) and MDA-MB-231 (**e**) cell apoptosis induced by indicated treatments (24 hours with 2-nM DNA origami structures or 12-nM peptides). Q1, percentage of cell necrosis; Q2, percentage of apoptosis-induced necrosis; Q3, percentage of cell apoptosis; Q4, percentage of alive cells. **b**, **d**, **f,** Percentage of apoptosis and apoptosis-induced necrosis (Q2 plus Q3) for MCF-7 (**b**), SK-BR-3 (**d**) and MDA-MB-231 (**f**) cells. Each point stands for the mean value of one biological replicate (n = 3). Each biological replicate includes 6 technical replicates. PBS stands for phosphate-buffered saline, and PEP stands for peptide alone. p* < 0.05. p** < 0.01. p*** < 0.001.

We also checked cleaved caspase-8 level, which is the key molecular indictor of apoptotic cascades^55^, by cell-based ELISA (Fig.6a). More cleaved caspase-8 was detected in MCF-7 cells showing a higher extent of apoptosis and apoptosis-induced necrosis (Fig.6b). Over all cell lines, the viability of cells treated by L6 was the lowest among different treatments (Fig.6c-e). This is the first time to circumvent the previously revealed TRAIL resistance of MCF-7 cell line. The half maximal inhibitory concentrations (IC_50_) of the peptide were significantly reduced after being patterned on DNA origami (W6, W9 and L6) (Extended Tab.1). Combining the data from these measurements and the GFP-DR5 clustering results showed clear correlations (Fig.6f-h), verifying that lower cell viabilities were most likely resulted from an effective DR5 clustering.

**Fig.6.**
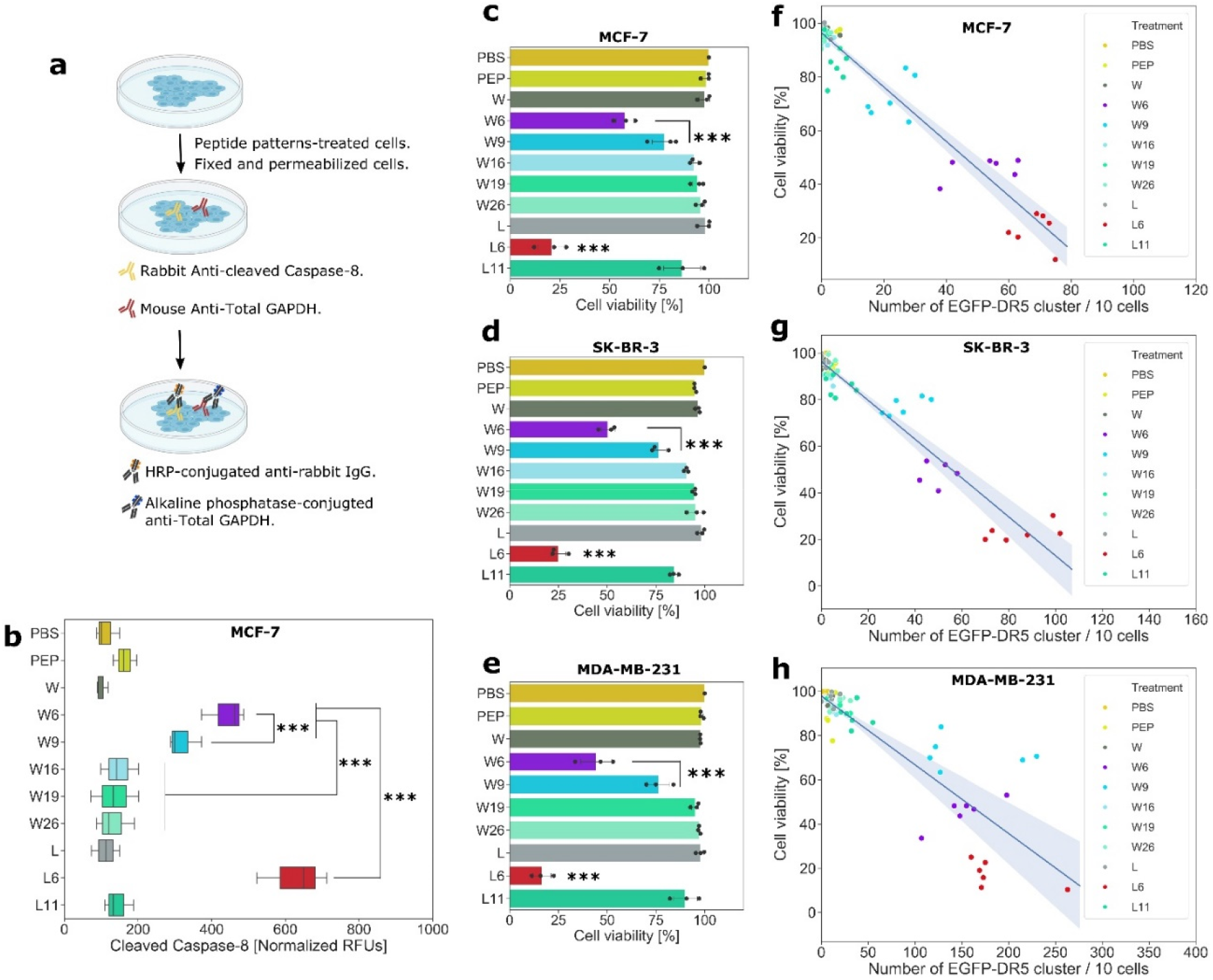
Cell viability. **a**, Workflow of cell-based ELISA for cleaved caspase-8 detection. **b**, Normalized (to GAPDH) cleaved caspase-8 level after the treatment (24 hours with 2-nM DNA origami structures or 12-nM peptides). **c**, **d**, **e**, Cell viability assay with indicated treatments (4 hours with 2-nM DNA origami structures or 12-nM peptides) from MCF-7 (**c**), SK-BR-3 (**d**) and MDA-MB-231 (**e**) cells. Each point stands for the mean value of one biological replicate (n = 3). Each biological replicate includes 6 technical replicates. **f**, **g**, **h**, Correlation of viability of MCF-7 (**f**), SK-BR-3 (**g**) and MDA-MB-231 (**h**) cells to the number (per 10 cells) of GFP-DR5 cluster. For each treatment, 6 groups of independent data are presented. PBS stands for phosphate-buffered saline, and PEP stands for peptide. p* < 0.05. p** < 0.01. p*** < 0.001.

## Conclusions

In conclusion, we demonstrated an effective strategy of using a DNA origami method to regulate death receptor clustering and following cell apoptosis. This offers a precise and reliable way to understand the importance of nanoscale ligand spatial organization and its control over apoptosis triggering. Notably, this approach does not rely on surface patterning and the stimulating patterns are displayed from solution. The fact that monomeric binders, as were used in this study, are able to trigger apoptosis could indicate that the pre-ligand clustering state of DR5 is less important than was previously thought and that the clustering of DR5, whatever the pre-ligand state may be, is enough to trigger apoptosis. By tuning the size of hexagonal TRAIL-mimicking peptide pattern, we conclude that inter-peptide distance for effective apoptosis was sub-10 nm. Surprisingly, this method worked on both TRAIL-sensitive and breast cancer cells that were previously deemed to be resistant. Our findings also reveal that precise spatial pattern screening of drug molecules at the nanoscale could be a potential way to alleviate some of the non-efficacy problems of currently approved TNFRSF-targeting drugs.

## Methods

### Peptide-DNA conjugate

We purchased the cyclic peptide (WDCLDNRIGRRQCVKL), with an azide modification at its C-terminal, from JPT Peptide Technologies. We purchased oligonucleotide (TAGATGGAGTGTGGTGTG), with a dibenzocyclooctyne (DBCO) modification at its 5-prime, from Integrated DNA Technologies. The molar ratio of peptide to oligonucleotide in the reaction was 10 to 1. We carried out the reaction in PBS pH 7.4, under room temperature, overnight. We used Amicon 3K filter tube (Millipore) to purify a small amount (100 uL, 100 nM) of conjugate. The process was done with centrifugation at 14,000 g for 30 min, and was repeated 3 times. We purified a big amount of conjugate (above 100 uL and 100 nM) via a proFIRE system (Dynamic Biosensors).

### Silver staining

We performed this using the ProteoSilver™ Silver Stain Kit (Sigma-Aldrich) according to its established protocol. Briefly, we placed gels in 100 ml of fixing buffer (50 ml Ethanol, 40 ml Milli-Q water, 10 ml Glacial Acid) overnight. Subsequently, we washed the gels for 10 min in 30% Ethanol solution, then in 200 ml of Milli-Q water for 10 min and sensitized the gels in 100 ml of sensitization solution (1 ml of Proteosilver sensitizer in 99 ml Milli-Q water). Following this, we washed the gels once more in Milli-Q water for 10 min, equilibrated the gels in silver solution (1 ml Proteosilver silver solution and 99 ml Milli-Q water) for 10 min, followed by two brief washes in Milli-Q water. The gels were then developed for approximately 5 min in 100 ml developer solution (5 ml of ProteoSilver Developer 1, 0.1 ml of ProteoSilver Developer 2 and 95 ml of Milli-Q water) after which the reaction was stopped by adding 5 ml of the provided stopping solution for at least 10 min. Stained gels were imaged using the GE LAS 4000 gel imager.

### Peptide-patterned DNA origami production

#### Step 1: p8064 scaffold DNA production

In a shaker at 37 °C, we cultured *Escherichia coli* strain JM109 in 250-mL 2× Yeast extract Tryptone growth medium (Sigma Aldrich) with 5 mM MgCl2 (Sigma Aldrich) until its optical density at 600 nm reached 0.5. Then we added the phage containing p8064 scaffold DNA to the bacteria at a multiplicity of infection of 1, after which the phage was amplified with shaking, under 37 °C, for 4 hours. We collected the culture, and then centrifuged at it 4,000 g for 30 min to pellet the bacteria. The supernatant, which contained the phage, was kept, followed by adding 10 g of PEG8000 (VWR international) and 7.5 g of NaCl (VWR international). We then incubated the supernatant with ice for 30 min and centrifuged it at 10,000 g for 30 min to pellet the phage. We re-suspended the phage in 10-mL Tris buffer (10 mM, pH 8.5, VWR international), and added 10 ml of a solution containing 0.2 M NaOH (VWR international) and 1% SDS. Then we denatured the phage protein coat by adding 7.5 ml of 3 M KOAc (VWR international), pH 5.5, and gently mixing, incubating on ice for 10 min. The sample was next centrifuged at 16,500g for 30 min to pellet the denatured phage proteins. The supernatant containing p8064 scaffold DNA was collected, and 50 ml 99.5% EtOH (Sigma Aldrich) was added, mixed gently and incubated on ice for another 30 min. The sample was centrifuged at 16,500g for 30 min pellet the p8064 scaffold DNA. The pellet was then washed with 75% EtOH and air dried at room temperature for 15 min, followed by re-suspending in 10 mM Tris, pH 8.5. The concentration and quality were characterized by UV-Vis (NanoDrop, Thermo Scientific) and a 2% agarose gel, respectively.

#### Step 2: DNA origami folding and purification

Staple oligonucleotides (Supplementary Table 1), with the concentration of 100 μM in Mili-Q water, were ordered from Integrated DNA Technologies in 96-well plates. DNA concentrations used for structure folding were as follows: 20 nM ssDNA scaffold, and 100 nM per staple DNA. For Cy5-labled DNA origami, 6 staple DNA strands of the structures were replaced by the same sequences but modified with Cy5 at their 5 primes. All wFS structures were folded in PBS by rapid heat denaturation (80 °C for 5 min) followed by cooling from 80 °C to 60 °C over 20 min, then from 60 °C to 24 °C over 14 hours. By using the same annealing program, all sFS structures were folded in the buffer containing 13 mM MgCl2 (Sigma-Aldrich), 5 mM TRIS (VWR international), and 1 mM EDTA (VWR international). Folded structures were purified and concentrated by using Amicon 100K filter tube (Millipore). The process included six-time washings with the folding buffer at 5,000 g for 2 min.

#### Step 3: Peptide attachment to DNA origami

The peptide-DNA conjugates were added with a tenfold excess to each protruding site on the DNA origami and incubated in the thermocycler (Bio-Rad) at 37 °C for 1 hour, followed by keeping under room temperature overnight.

#### Step 4: Removal of excess conjugate

This was carried out also by using Amicon 100K filter tubes (Millipore). The removal process included six-time washings with the folding buffer at 5,000g for 2 min. The final concentrations of peptide-patterned DNA origami were measured at UV-Vis A260 on Nanodrop (Thermo Scientific).

### Agarose gels electrophoresis

2% agarose (Sigma-Aldrich) gels were cast in 0.5× TBE buffer (VWR) supplemented with 10 mM MgCl2 and 0.5 mg/mL ethidium bromide (Sigma-Aldrich). For all samples, gels were run in 0.5× TBE buffer supplemented with 10 mM MgCl2 at 90 volts for 3 hours on ice. Gels were imaged under a GE LAS 4000 imager.

### Gel-based peptide quantification of DNA origami

#### Step 1: DNA origami digestion

Under 37 °C, 10 μL of 20-nM DNA origami structures (with different number of peptide attachment) in PBS were incubated with 0.1U/μL DNase I (Invitrogen) and 2mM MgCl2 for 25 min. The goal of this process was to completely degrade DNA origami template, releasing peptides from the structure.

#### Step 2: gel electrophoresis and peptide staining

The samples were then run on 4-20% gradient PAGE gel (Bio-Rad) with 1-L buffer containing 3 g of Tris base (VWR international), 14.4 g of glycine (VWR international) and 1 g of SDS (Sigma Aldrich). After the running, the gel was stained by the method of silver staining.

#### Step 3: Peptide band intensity quantification

In Image J, the band areas (same size for all bands on one gel) were selected, and the mean pixel intensities of the areas were used to compare the peptide amount.

#### Step 3: Peptide occupancy rate calculation

We calculated the peptide amount of each sample based on their band intensity on the polyacrylamide gels: Peptide amount of each sample = (Known peptide amount of the peptide only sample × band intensity of each sample) / band intensity of the peptide only sample. We then calculated the protrusion site occupation (by peptide) percentage with the equation: Occupation percentage = peptide amount of each sample / (DNA origami amount in each sample × the number of protrusion sites) × 100%.

### Atomic force microscopy

10 μL of samples, with the concentration of 1.5 nM in imaging buffer [10 mM MgCl2 (Sigma-Aldrich), 5 mM TRIS (VWR), and 1 mM EDTA (VWR)] were dropped to freshly cleaved mica for 30-second incubation. 4 μL of NiSO4 (5 mM, VWR) was added for further 5-min incubation. We then washed the sample-micasurface by 1-mL imaging buffer. We used a cantilever AC40 (Bruker) with a nominal spring constant of 0.09 N/m to carry out the AFM (JPK instruments Nanowizard 3 ultra) imaging.

### DNA PAINT

#### Step 1: Sample preparation for DNA-PAINT

Glass microscope slides (VWR) and coverslips (1.5H, VWR) were cleaned with acetone and isopropanol before drying. Double sided scotch tape was placed onto the slides in two parallel stripes approximately 0.8 cm apart and the clean coverslips were placed on it to create flow chambers as described before (ref.). The channel was flushed with 1mg/mL of biotinylated-BSA (Sigma Aldrich) in Buffer A (10mM Tris-HCl, 100mM NaCl, 0.05% Tween-20, pH 7.5) and incubated for 2 minutes. The channel was then washed with Buffer A. The channel was then flushed with 0.5mg/mL streptavidine (Thermo Scientific) in Buffer A and incubated for 2 minutes. The channel was washed then with Buffer A. After this 80nm AuNP solution (Sigma Aldrich), used as fiducial markers, re-suspended in Buffer A was flushed into the channel and incubated for 2min followed by a washing step with Buffer A. The channel was then washed with Buffer B (5mM Tris-HCl, 10mM MgCl2, 1mM EDTA, 0.05% Tween-20, pH 8) before structures carrying TRAIL-peptides with DNA-PAINT docking sites and six biotin sites for immobilization resuspended in Buffer B were flushed in at 200pM concentration and incubated for 5 minutes, followed by washing with Buffer B. The channel was then flushed with imaging buffer and sealed with epoxy glue. The imaging buffer used was based on Buffer B and contained oxygen scavengers 2.4mM PCA (Sigma Aldrich) and 10nM PCD (Sigma Aldrich) and 1mM Trolox (Sigma Aldrich) at previously along with Atto-550 labelled imager strands (IDT) at the concentration of 10nM.

The samples were imaged with a microscope with a Nikon Eclipse Ti-E microscope frame with the Perfect Focus system (Nikon Instruments) and an objective-type TIRF configuration using an iLAS2 circular TIRF module (Gataca systems). For magnification a 1.49 NA CFI Plan Apo TIRF 100× Oil immersion objective (Nikon Instruments) was used with a 1.5x auxiliary Optovar magnification resulting in a final pixel size of 87nm. For illumination an OBIS 561nm LS 150mW laser (Coherent) was used with custom iLas input beam expansion optics (Cairn) optimized for reduced field super resolution imaging. The laser beam was passed through before the objective a filter cube (89901, Chroma Technology) containing an excitation quadband filter (ZET405/488/561/640x, Chroma Technology), a quadband dicroic (ZET405/488/561/640bs, Chroma Technology) and a quadband emission filter (ZET405/488/561/640m, Chroma Technology). The collected light was spectrally filtered with an additional emission filter (ET595/50m, Chroma Technology) before entering the iXon Ultra 888 EMCCD camera (Andor) used for recording. The Micromanager software were used for acquiring 12000 frames long time lapses of samples using frame-transfer mode of the camera, 300 ms exposure time, 10 MHz readout rate and no EM gain.

#### Step 2: Pre-processing of DNA-PAINT data

Localization coordinates in the collected time-lapses were calculated using the Picasso Localize program from the Picasso sofware package (ref) using 2000 for minimum net gradient for localization identification and MLE algorithm with 0,001 and 1000 set as convergence criterion and maximum number of allowed iterations respectively. The localizations were then drift-corrected using the RCC algorithm with 200 frame fragment size in the Picasso render program and localizations belonging to the gold nanoparticles used as fiducial markers were selected and exported using the “export selected localizations” feature of the program for later filtering. Following this the low precision localization (localization precision > 0.03 camera pixels), asymmetric localizations (localization ellipticity < 0.1) and multi-localizations (photon count > mean photon count + 2 STD) were removed along localization belonging to the gold nanoparticles using a custom script (“Loc-prec-filtering”). Finally, the localizations belonging to single structures were selected using the “pick similar” feature of the Picasso Render software (circular pick regions, 1.5 camera pixel diameter, pick similar range of 2.0 std) starting from 30 manually picked structures and the localization were undrifted a second time using the software’s “undrift from picked” feature

### Processing of DNA-PAINT images of peptide-patterned DNA origamis

#### Detection of DNA origami structure in localization data

DNA origami structures were detected using a custom script from the cleaned localization data. The localizations were rendered into a low resolution image (20x oversampling) and clusters of localizations were detected using contour detection and sorted into origami ROIs centered around the detected clusters. ROIs containing noise due to unspecific binding of imagers were removed based on the cluster size and temporal span of localization within the ROI.

#### Quantification of TRAIL peptides in structure ROIs

TRAIL proteins were detected using a custom script in the localizations sorted into origami ROIs. Localizations in each origami ROI were rendered into a high resolution images (60x, 60x, 100x, 120X, 120X and 150x oversampling for the 37nm, 28nm, 19nm, 16nm, 9nm and 5nm origami structures), the intensity of images was normalized and a gaussian smoothing filter was applied to them. Local maxima in these images were detected and maxima closer than ~0.5 x the designed site distance were merged. To only keep maxima belonging to a single origami structure a euclidian distance based clustering was applied to the maxima and only the most central clusters with the highest number of maxima were kept. The coordinates of these maxima were then exported as detected positions of TRAIL peptides. For quantification the TRAIL peptide per origami distribution were calculated from counting the detected TRAIL peptides in each origami ROI and for the mean nearest neighbour distance distribution the distance to the closest neighbour was calculated for each TRAIL position detected in an origami ROI and the mean was calculated from that for each origami ROI in each data set.

### Cell culture

Cell lines were purchased from ATCC. All of them were cultured in Dulbecco’s Modified Eagle’s Medium (DMEM) (Sigma-Aldrich) with 20% heat-inactivated FBS (Gibco), 100 U/mL penicillin-streptomycin (Gibco) in a humidified environment containing 5% CO_2_ at 37 °C. To establish GFP-DR5-expressing cell lines, we used Lipofectamine 3000 reagent (Thermo Fisher Scientific) to transfect GFP-tagged human tumor necrosis factor receptor superfamily 10b (GFP-TNFRSF10B/GFP-DR5) plasmid (OriGene) into the cell lines. For each well of a 6-well plate, cells were cultured to be 70-90% confluent for transfection. Then 2.5 ug of GFP-TNFRSF10B plasmid was mixed with 3.75 μL of Lipofectamine 3000 reagent and 5 μL of P3000 reagent (reagent in the kit) for 15-min incubation. Then we added the DNA-lipid complex to each cell well for 2 days. The transfected cells were then used immediately for the following treatment experiments.

### Confocal data collection and image analysis

1×10^6^ cells per well were cultured on a coverslip in each well of a 6-well plate 24 h prior to the transfection of GFP-DR5 plasmid. After 2 days, cells were washed with fresh DMEM medium containing 20% heat-inactivated FBS and 100U/mL penicillin-streptomycin. Then the cells were treated with peptide-patterned DNA origami structures (Cy5-free or Cy5-labeled), at certain concentrations and for certain periods (as indicated in corresponding Figures or the caption of Figures). Finally, cells were fixed in 4% paraformaldehyde for 15 min, washed, and stained by Fluoroshield Mounting Medium with DAPI (Abcam). Cells were imaged on LSM710 (Zeiss). For GFP-DR5 cluster quantification in ImageJ, images were converted into 8-bit images, and thresholded for the next processing. The Analyze Particles plug-in for ImageJ, as an automatic particle counting tool, was used to count the number of GFP-DR5 clusters in each image. For colocalization analysis between Cy5-labeled DNA origami and GFP-DR5 clusters, the analysis was performed with Colocalization Analysis plug-in in ImageJ.

### Flow cytometry

In a 6-well plate, 1×10^7^ cells per well were cultured for 24 h prior to treatments. Cells were then treated with 2-nM peptide-patterned DNA origami (equals to 12 nM peptide) for 24 h. All cells (including dead and detached cells that were present in the medium) were collected (centrifuge at 3000 RPM for 5 min) and washed with cold PBS for 3 times. Cells were resuspended, and then stained with annexin V-FITC and PI, sequentially, according to the commercial protocol of the Dead Cell Apoptosis Kit with Annexin V-FITC and PI (Thermo Fisher Scientific).

### Cleaved caspase-8 detection

We used the method of Cell-based ELISA with using a Human Cleaved Caspase-8 (Asp391) Immunoassay kit (R&D systems).

#### Step 1: culture, treat, fix and block cells

1.5×10^4^ cells in 100 μL of DMEM medium per well were seeded in a 96-well black polystyrene microplate with clear bottom (Corning) and cultured overnight. Cells were then treated with 2-nM peptide-patterned DNA origami (equals to 12 nM peptide) for 4 h. Cells were then fixed by replacing the medium with 100 μL of 4% formaldehyde in PBS, for 20 min under room temperature. After the fixation, cells were washed by the Wash Buffer, kept in 100 μL of the Quenching Buffer for 20 min, again washed by the Wash Buffer, and kept in 100 μL of Blocking Buffer for 1 hour.

#### Step 2: incubation of primary and secondary antibodies

After removing the Blocking Buffer and washing the cells with Wash Buffer, 100 μL of the primary antibody mixture containing rabbit anti-cleaved caspase-8 (Asp391) and mouse anti-total GAPDH was added for 16-hour incubation under 4 °C. The primary antibodies were removed, and the cells were washed by Wash Buffer. Then 100 μL of the secondary antibody mixture containing HRP-conjugated anti-rabbit IgG and AP-conjugated anti-mouse IgG was added for 2-hour incubation under room temperature to bind targeting primary antibodies.

#### Step 3: Fluorogenic detection

The secondary antibodies were removed, and the cells were washed with Wash Buffer. 75 μL of substrate of HRP was added for 30 min incubation at room temperature, followed by adding 75 μL of substrate of AP for an additional 30 min incubation. The signals were then read under a multimode microplate reader (Varioskan™ LUX): with excitation at 540 nm and emission at 600 nm for Cleaved Caspase-8 (Asp391) detection; with excitation at 360 nm and emission at 450 nm for total GAPDH detection.

### Cell viability assay

We used the method of ATP-based luminescent cell viability assay with the Kit named CellTiter-Glo Luminescent Cell Viability Assays (Promega). 5×10^4^ cells in 100 μL of DMEM medium per well were seeded in a 96-well opaque white polystyrene microplate (Corning) and cultured for 24 hours. Cells were treated with various concentrations of peptide-patterned DNA origami structures (as indicated in corresponding Figures or the caption of Figures). After 48-hour incubation, the plate was taken out from the cell incubator, and equilibrated at room temperature for 30 min. 100 μL of CellTiter-Glo reagent (Promega) was added to each well, followed by mixing for 2 min on an orbital shaker to induce cell lysis. The plate was then incubated at room temperature for 10 min to stabilize the luminescent signal. Finally, the luminescence was recorded on a multimode microplate reader (Varioskan™ LUX). The results of control wells containing medium without cells were used as the background luminescence. % viable cells = (Luminescence sample - Luminescence background) / (Luminescence PBS – Luminescence background) × 100.

### Statistical analysis

All experiments were performed in multiple distinct replicates, as indicated in the text and figure legends. Statistical analysis was performed using Graphpad Prism 7.0 (GraphPad). Data were analyzed using two-tailed Student’s t-tests for 2 groups and one-way ANOVA followed by Tukey post tests for multiple groups. Results are expressed as mean ± S.D. unless otherwise indicated. For each box-and-whisker plot, center line is the median and whiskers represent the minimum and maximum values.

## Acknowledgements

We would like to acknowledge support from the Knut and Alice Wallenberg Foundation for B. Högberg (KAW 2017.0114 and KAW 2017.0276) and from the European Research Council ERC for B. Högberg (Acronym: CellTrack GA# 724872).

## Author contributions

Y.W., I.B. and B.H. conceived the study and designed experiments. Y.W. performed most of the experimental work on origami, cellular imaging and data analysis. I.B. performed a significant proportion of the experimental work on origami, gel electrophoresis and cellular experiments. F.F. performed the DNA-PAINT data collection and analysis. All the authors contributed to writing the manuscript.

## Competing interests

The authors declare no competing interests.

---

Extended Figures and methods for

| **Index** | **Contents** |
| --- | --- |
| **Extended Fig.1** | Design of L-structure |
| **Extended Fig.2** | Computational 3D structure of L designs |
| **Extended Fig.3** | Agarose gel electrophoresis of W and L |
| **Extended Fig.4** | Basic structural information of the peptide and TRAIL trimer |
| **Extended Fig.5** | Scheme of peptide-DNA conjugation |
| **Extended Fig.6** | Conjugates gel electrophoresis followed by silver staining |
| **Extended Fig.7** | Current interpretation of apoptosis-driving hexagonal DR5 clusters |
| **Extended Fig.8** | Illustration on pattern designs on W |
| **Extended Fig.9** | Design of the template for L6 |
| **Extended Fig.10** | Design of the template for L11 |
| **Extended Fig.11** | Nucleotide-to-nucleotide distance measurement through oxDNA |
| **Extended Fig.12** | Agarose gel electrophoresis of peptide-attached W structures |
| **Extended Fig.13** | Agarose gel electrophoresis of peptide-attached L structures |
| **Extended Fig.14** | Pre-processing of DNA-PAINT data |
| **Extended Fig.15** | Plots of localization properties and their respective cutoffs for filtering |
| **Extended Fig.16** | Detection of DNA origami ROI coordinates from DNA-PAINT data |
| **Extended Fig.17** | Detection of peptides in origami ROIs extracted from DNA-PAINT data |
| **Extended Fig.18** | DNA-PAINT images of W37 structures |
| **Extended Fig.19** | DNA-PAINT images of W26 structures |
| **Extended Fig.20** | DNA-PAINT images of W19 structures |
| **Extended Fig.21** | DNA-PAINT images of W16 structures |
| **Extended Fig.22** | DNA-PAINT images of W9 structures |
| **Extended Fig.23** | DNA-PAINT images of W6 structures |
| **Extended Fig.24** | Protruding DNA site occupancy from DNA-PAINT |
| **Extended Fig.25** | inter-peptide distance distribution from DNA-PAINT |
| **Extended Fig.26** | Gel-based peptide quantification of L6 |
| **Extended Fig.27** | DR5 clusters induced by L6 attached with different number of peptides |
| **Extended Tab.1** | The half maximal inhibitory concentration ($IC_{50}$) of peptide |

**Extended Fig.1.**
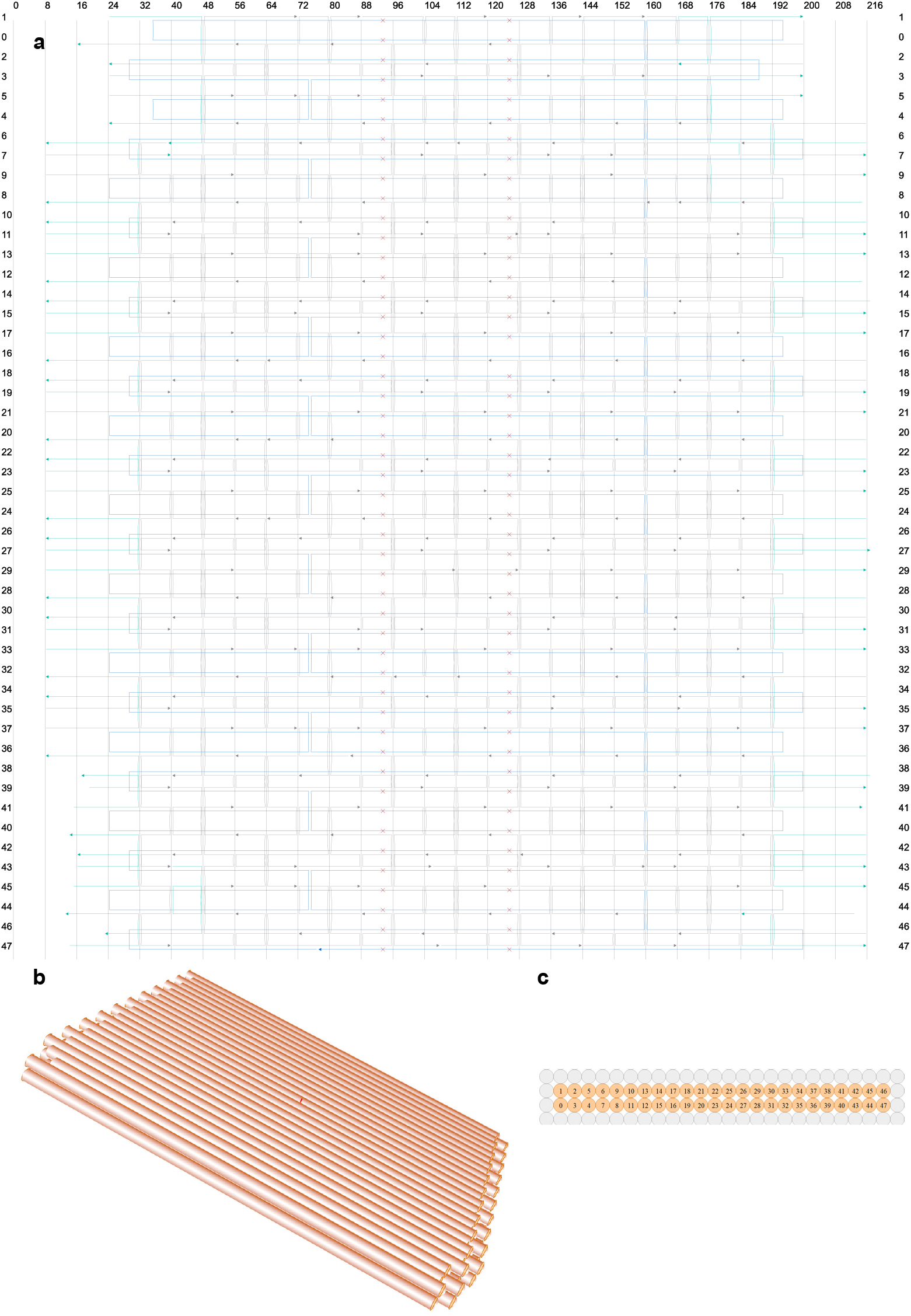
Design of L. **a,** the design blueprint of L in cadnanoSQ. The scaffold DNA is in blue, while DNA staples are in gray. The DNA staples colored in cyan have polyA sequences at their non-hybridized parts, which can help reducing aggregation. **b,** 3D view of L. **c,** the cross-section view of L in cadnanoSQ.

**Extended Fig.2.**
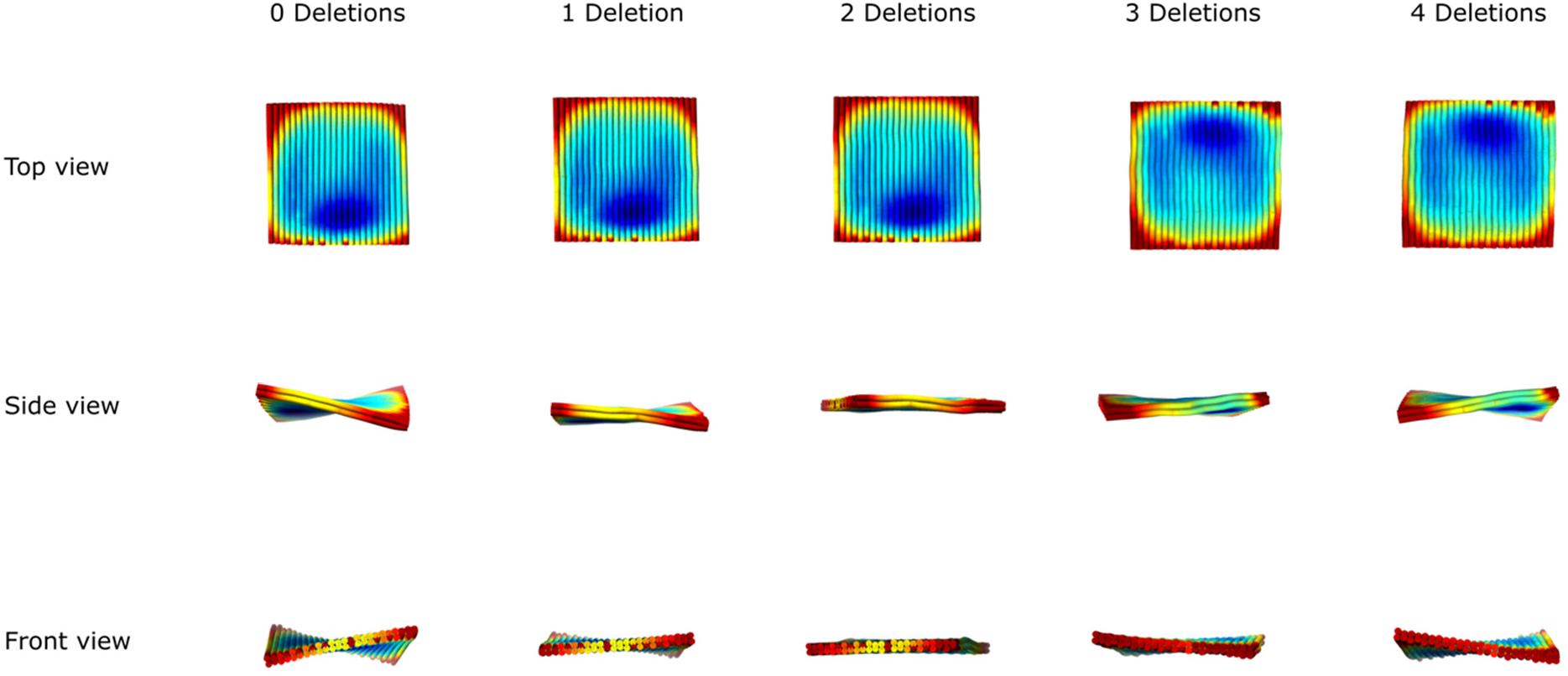
The computational feedback on the 3D structure of different L designs. As shown in Extended Fig.1, there are 2 columns of deleted bases, where the red “×” marks locate, in the scaffold DNA routing. We chose the number of columns for bases deletions by checking the 3D structure prediction in CanDo (https://candodna-origami.org/). The template with 2 columns of base deletions (2 Deletions) shows a relatively flat 3D configuration, while other designs (0 Deletions, 1 Deletions, 3 Deletions and 4 Deletions) present twisted configurations. Thus, we used the template having 2 Deletions for next peptide patterning.

**Extended Fig.3.**
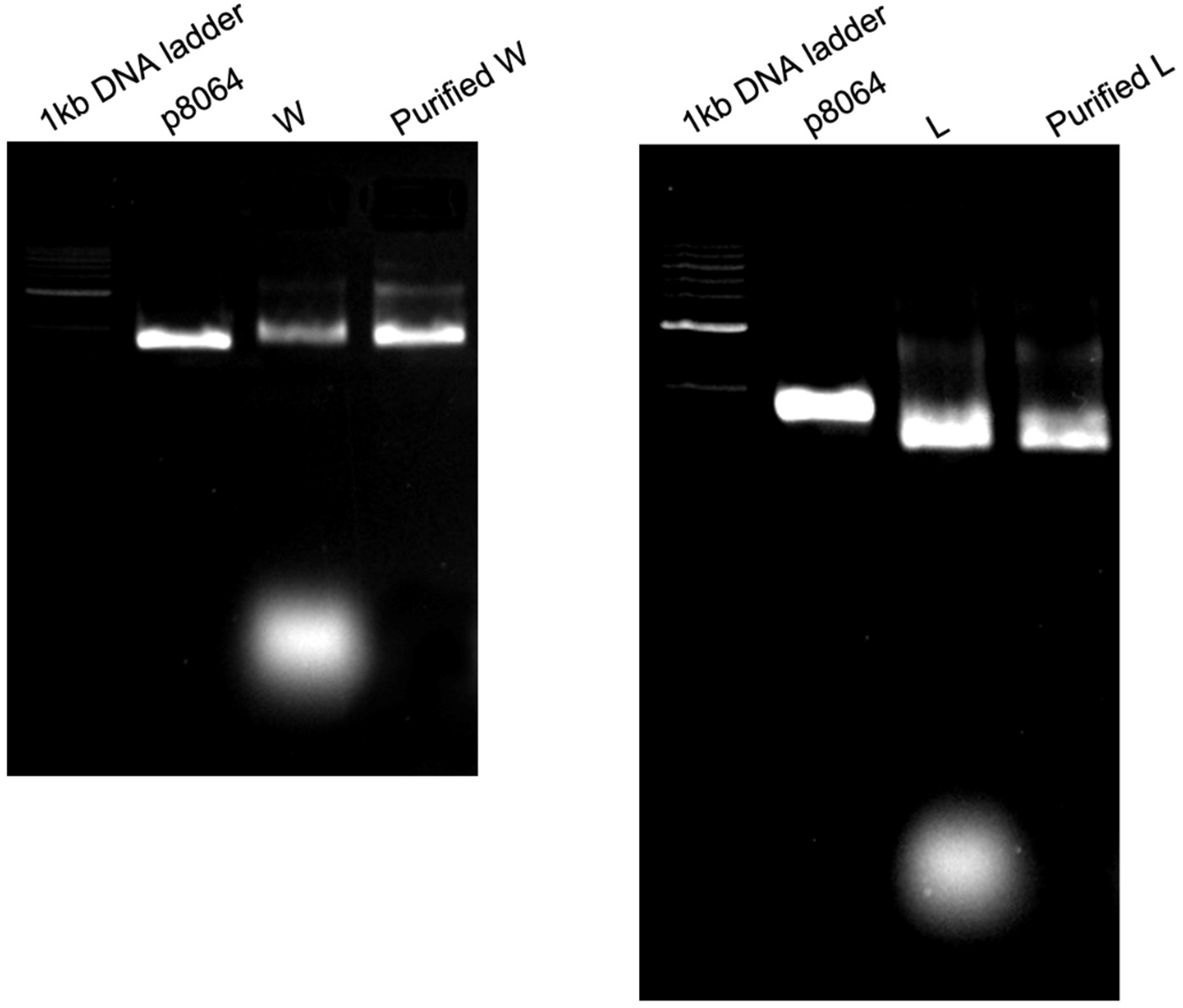
2% agarose gel electrophoresis of W (Left) and L (Right), before and after purification. The band (the bottom one) stands for the DNA staples, which were completely washed away by the purification process.

**Extended Fig.4.**
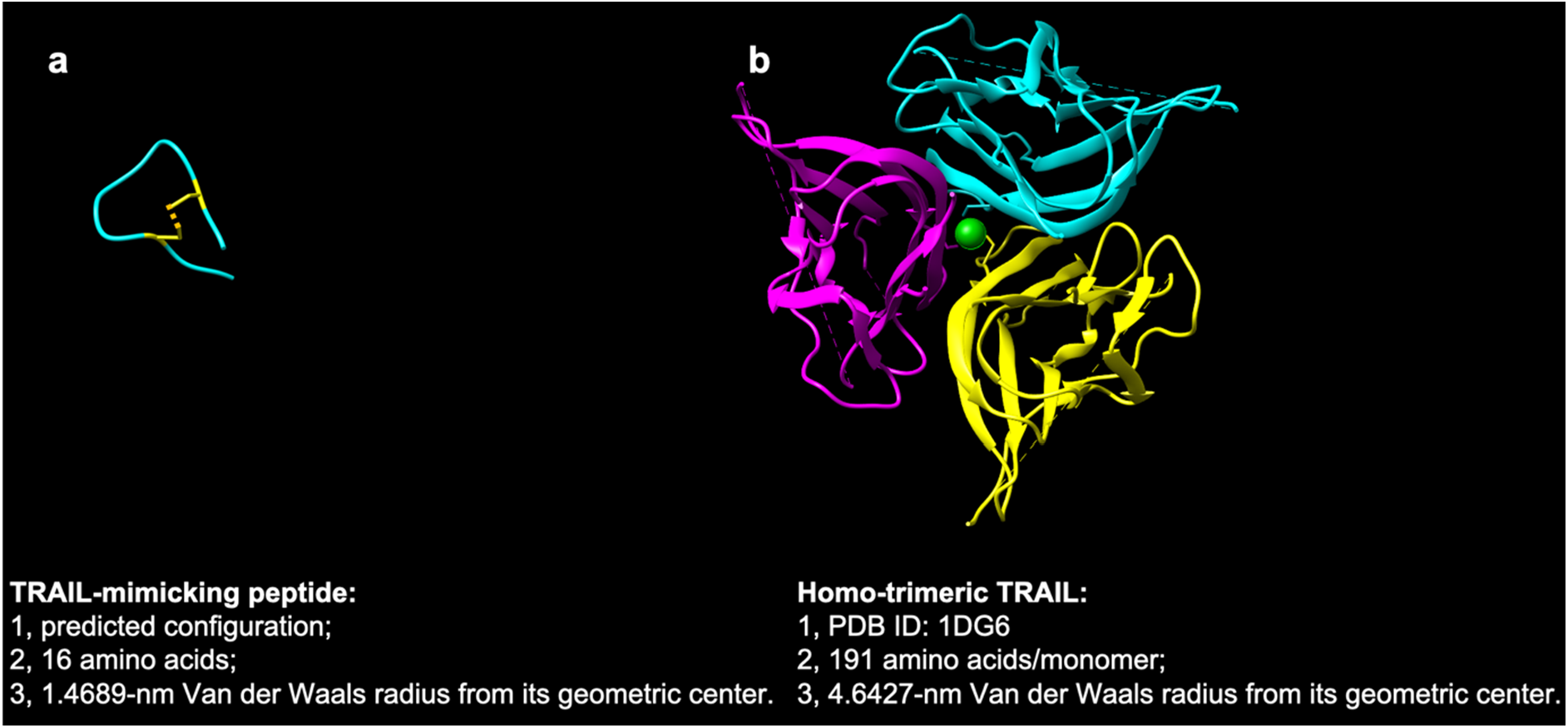
Basic structural information of the peptide and TRAIL trimer. **a**, peptide; **b**: homo-trimeric TRAIL. To predict the spatial structure of the peptide, its sequence (WDCLDNRIGRRQCVKL) was submitted to PEP-FOLD server (https://bioserv.rpbs.univ-paris-diderot.fr/services/PEP-FOLD/) to compute its PDB coordinates information, which was then visualized and analyzed in Chimera (https://www.cgl.ucsf.edu/chimera/) and YASARA (http://www.yasara.org/). The peptide is cyclized by a disulfide bond between two cysteines. The Human TRAIL trimer structure, with 1DG6 as its PDB ID, was directly sourced from RCSB PDB (https://www.rcsb.org/). Its parameter of Van der Walls radius from its geometric center was also analyzed in YASARA.

**Extended Fig.5.**
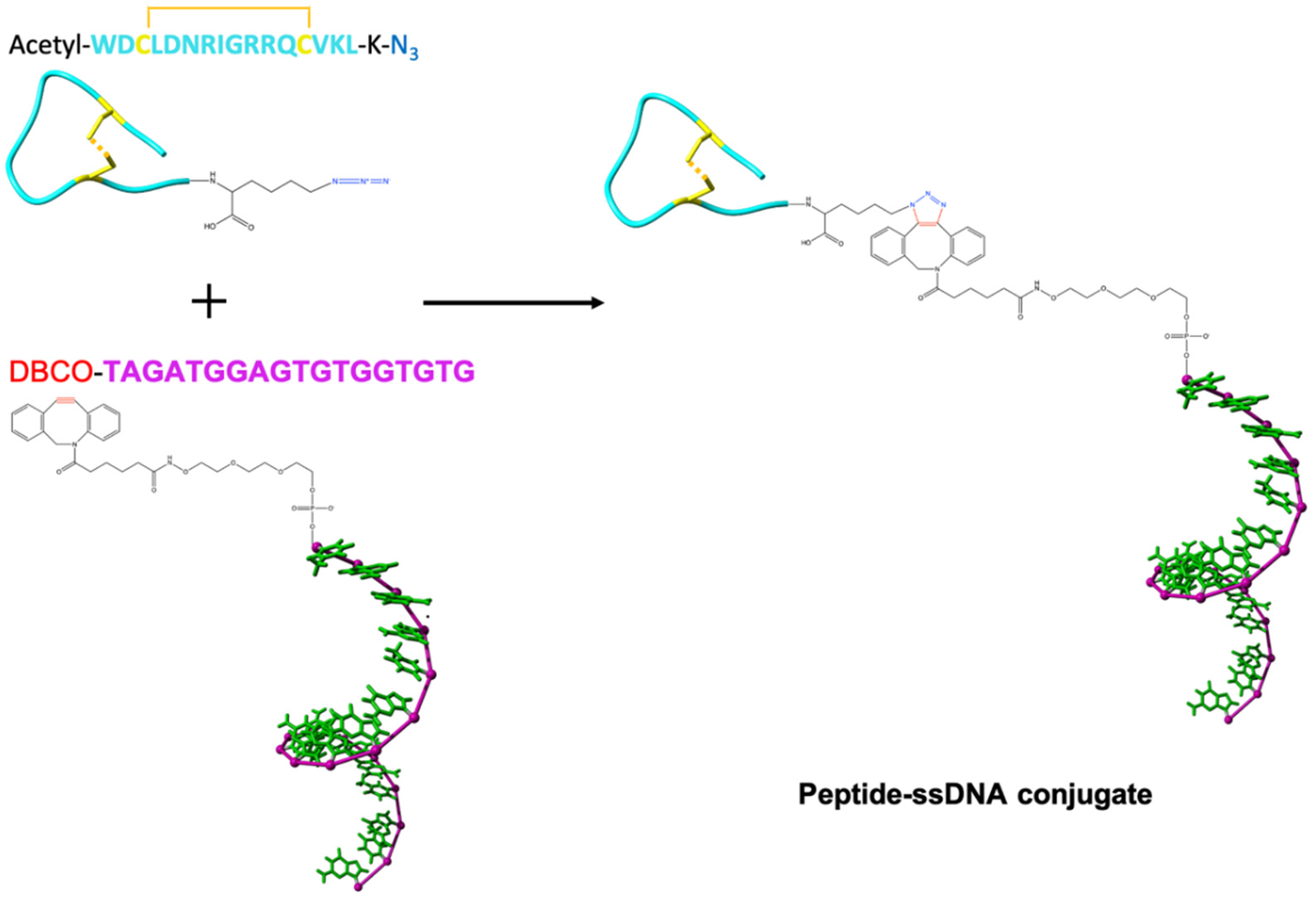
Peptide-DNA conjugation. The peptide is modified with an azide group at its C-terminal; the 18-base long ssDNA is tagged a DBCO at its 5-prime. The reaction is a copper free click chemistry reaction, which has a high reaction efficiency in PBS (pH 7.4), under room temperature.

**Extended Fig.6.**
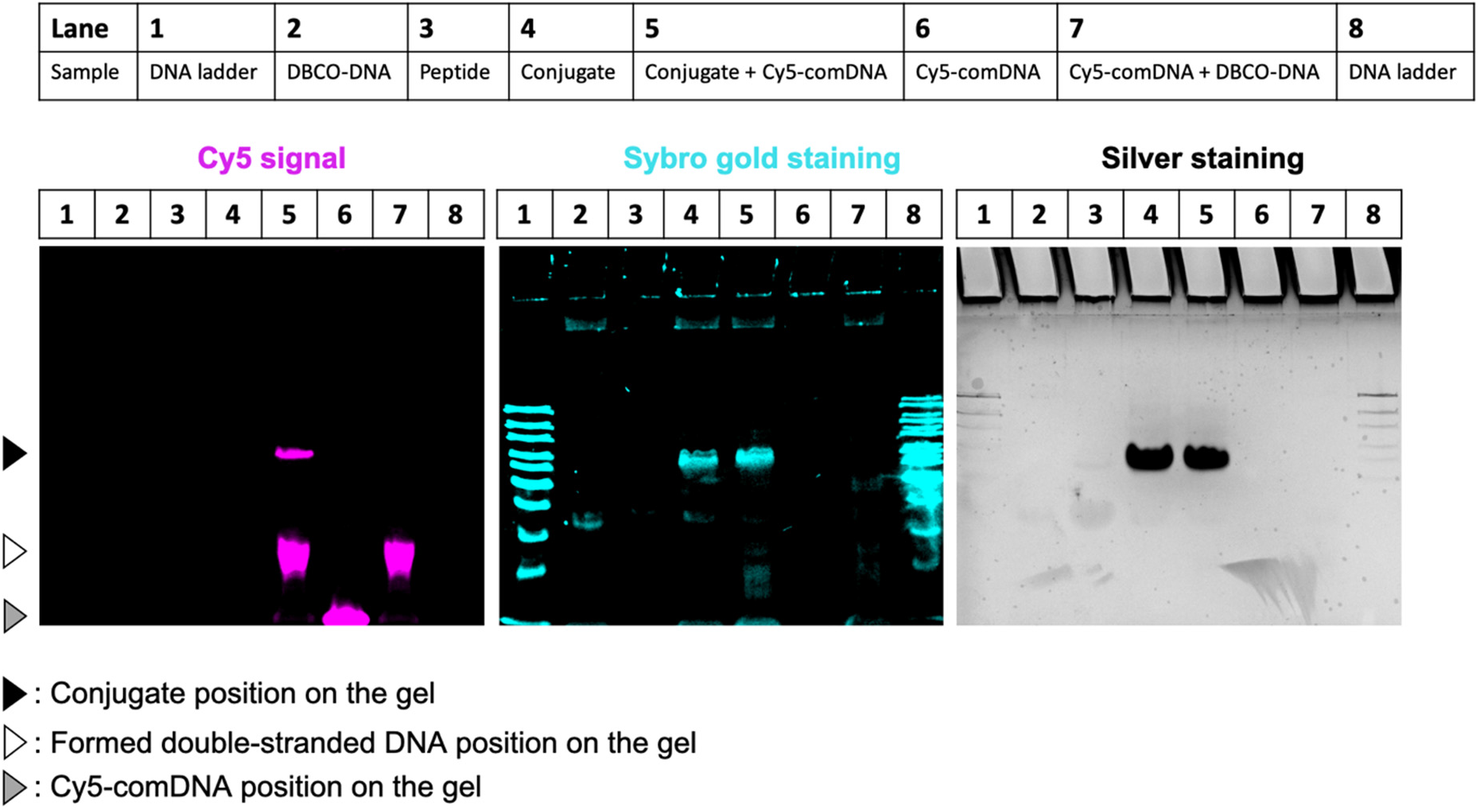
Validation the peptide-DNA conjugation by gel followed by silver staining. 18% native polyacrylamide gel was used. We used a Cy5-labled complementary DNA (Cy5-comDNA) to hybridize the DNA of the conjugate, thus we can visualize the conjugate under Cy5 channel. All of the samples are indicated as in the figure. The gel was imaged under Cy5 channel; nucleic acids on the gel was stained by SYBR gold; peptides were stained by silver staining.

**Extended Fig.7.**
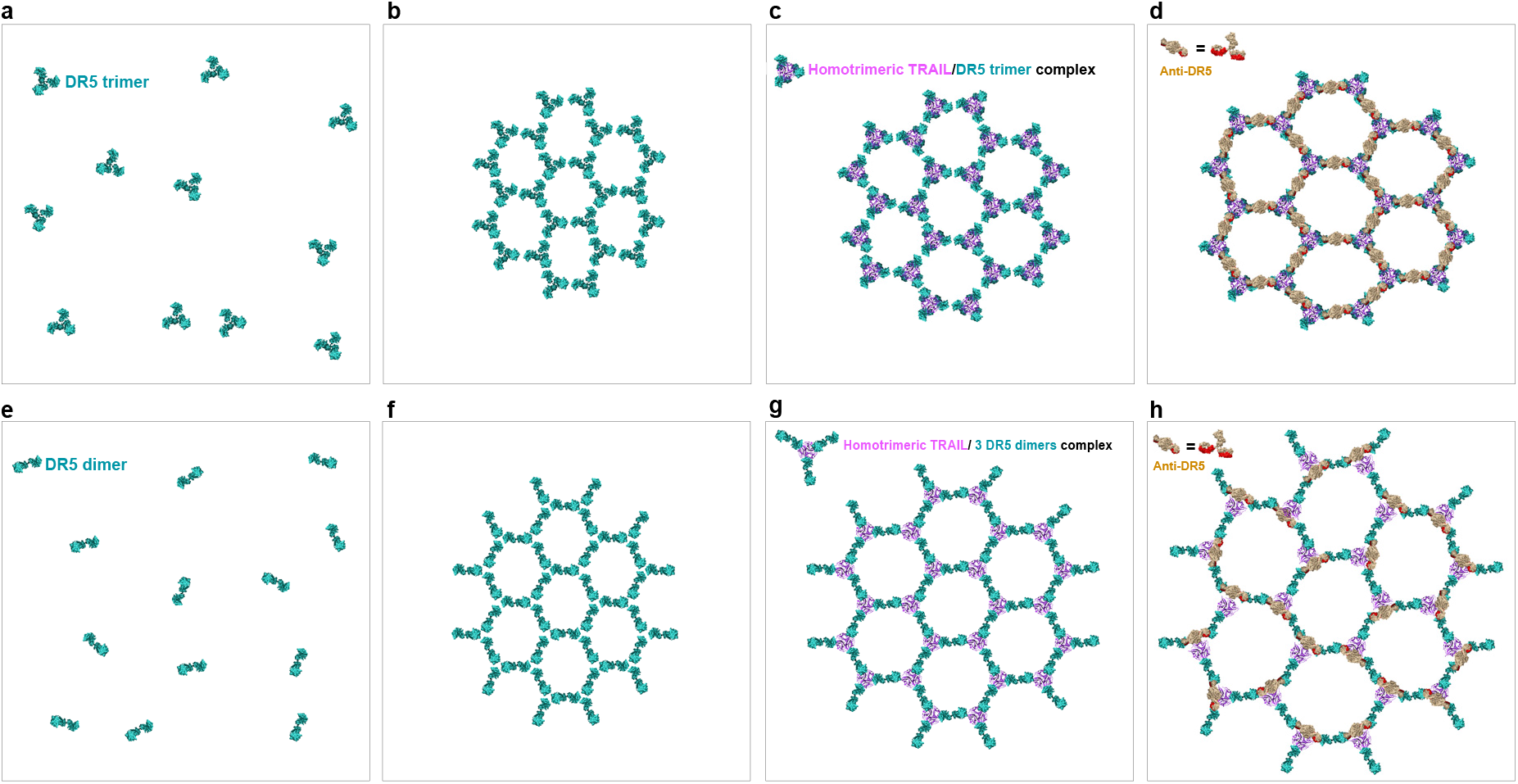
Current interpretation of apoptosis-driving hexagonal DR5 networks (top view from the outside of a cell surface). **a,** Surface representations of pre-ligand DR5 trimers. **b,** The proposed active hexagonal DR5 network via dimerization of DR5 trimers. **c,** The proposed active hexagonal network of homotrimeric TRAIL/DR5 trimer complexes. **d,** The hexagonal homo-trimeric TRAIL/DR5 trimer network promoted and maintained by the antibody AMG 655. **e,** Surface representations of pre-ligand DR5 dimers. **f,** The proposed active hexagonal DR5 network via trimerization of DR5 dimers. **g,** The proposed active hexagonal network of homo-trimeric TRAIL/DR5 dimer complexes. **h,** The hexagonal homo-trimeric TRAIL/DR5 dimer network promoted and maintained by antibody AMG 655.

**Extended Fig.8.**
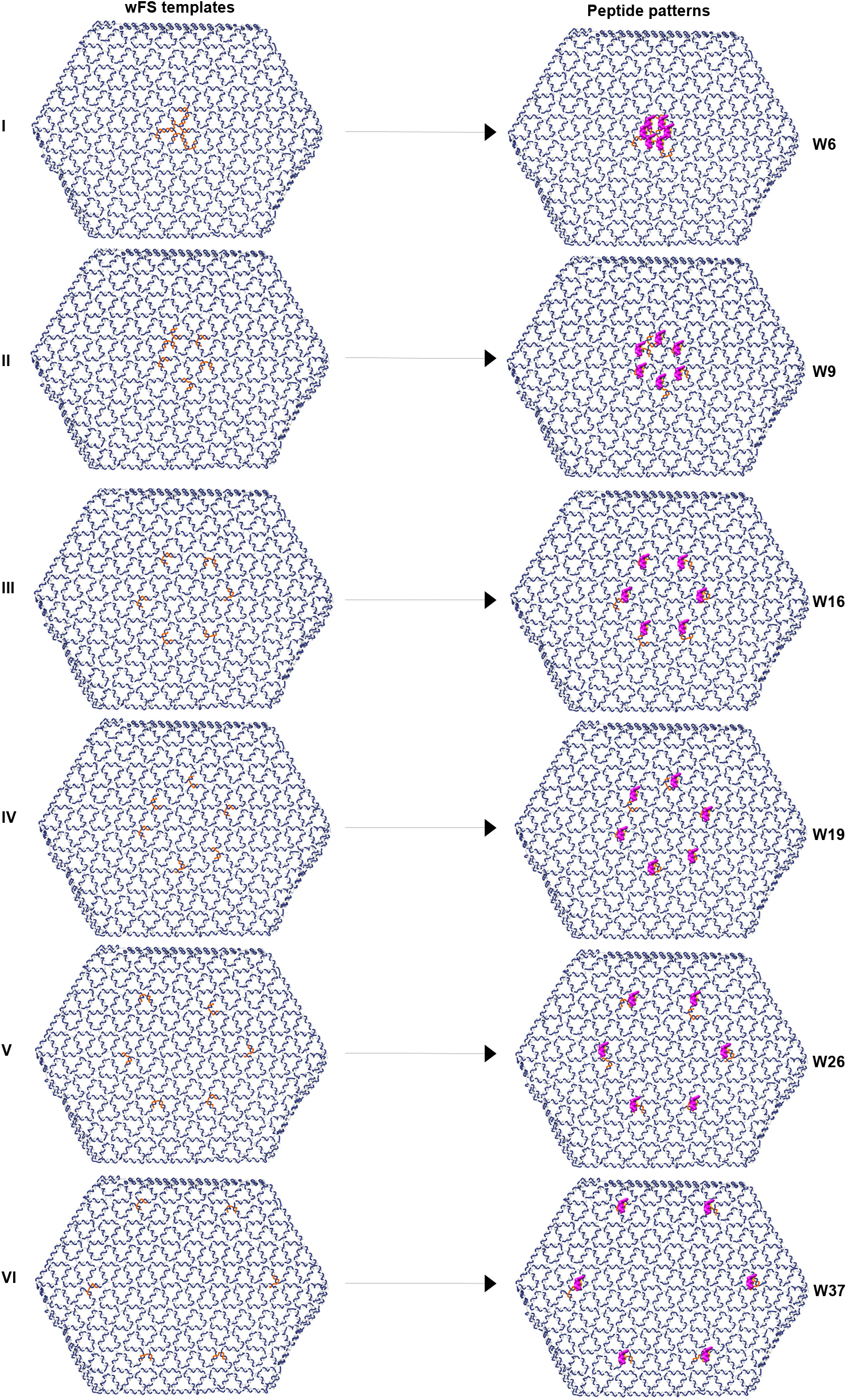
Designs of peptide patterns on W. The sizes of hexagonal patterns are decided by the choices of specific DNA staples on W. These staples were colored in orange, while blue is the scaffold DNA and gray means other DNA staples. These staples in orange contain protruding single-stranded DNA at their 5 primes, thus they can hybridize with the DNA-peptide conjugate.

**Extended Fig.9.**
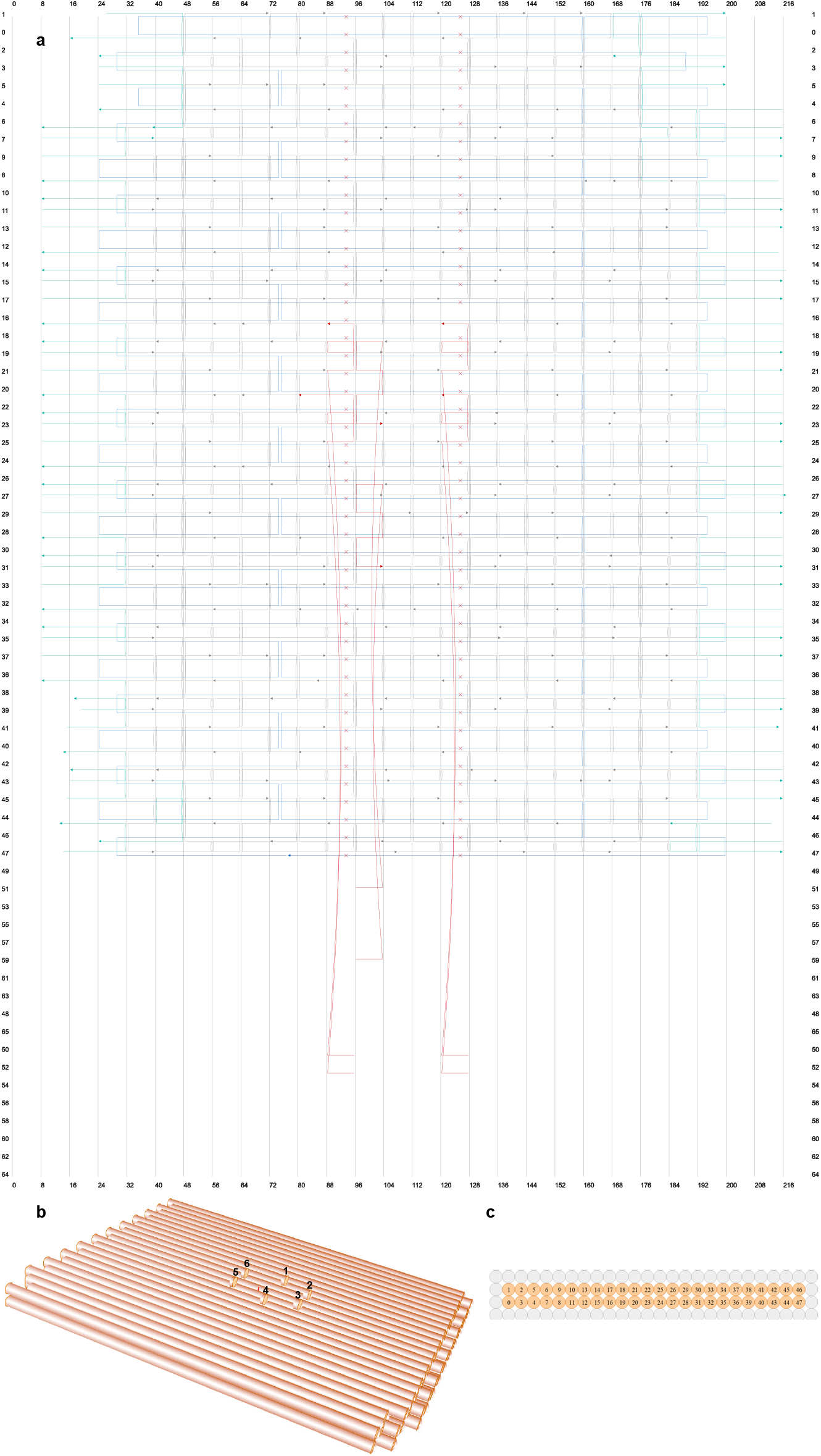
Design of the template for L6. **a,** the design blueprint of the template in cadnanoSQ. The scaffold DNA is in blue, while DNA staples are in gray. The DNA staples colored in cyan have polyA sequences at their non-hybridized parts, which can help reducing aggregation. The 6 staples having protruding sequences at their 5 primes, for the docking of DNA-peptide conjugate, were colored in red. **b,** the 3D view of the tempalte, showing that the 6 protruding sites are presenting a hexagonal pattern, and the distance between adjacent sites are around 5 nm. **c,** the cross-section view of the template in cadnanoSQ.

**Extended Fig.10.**
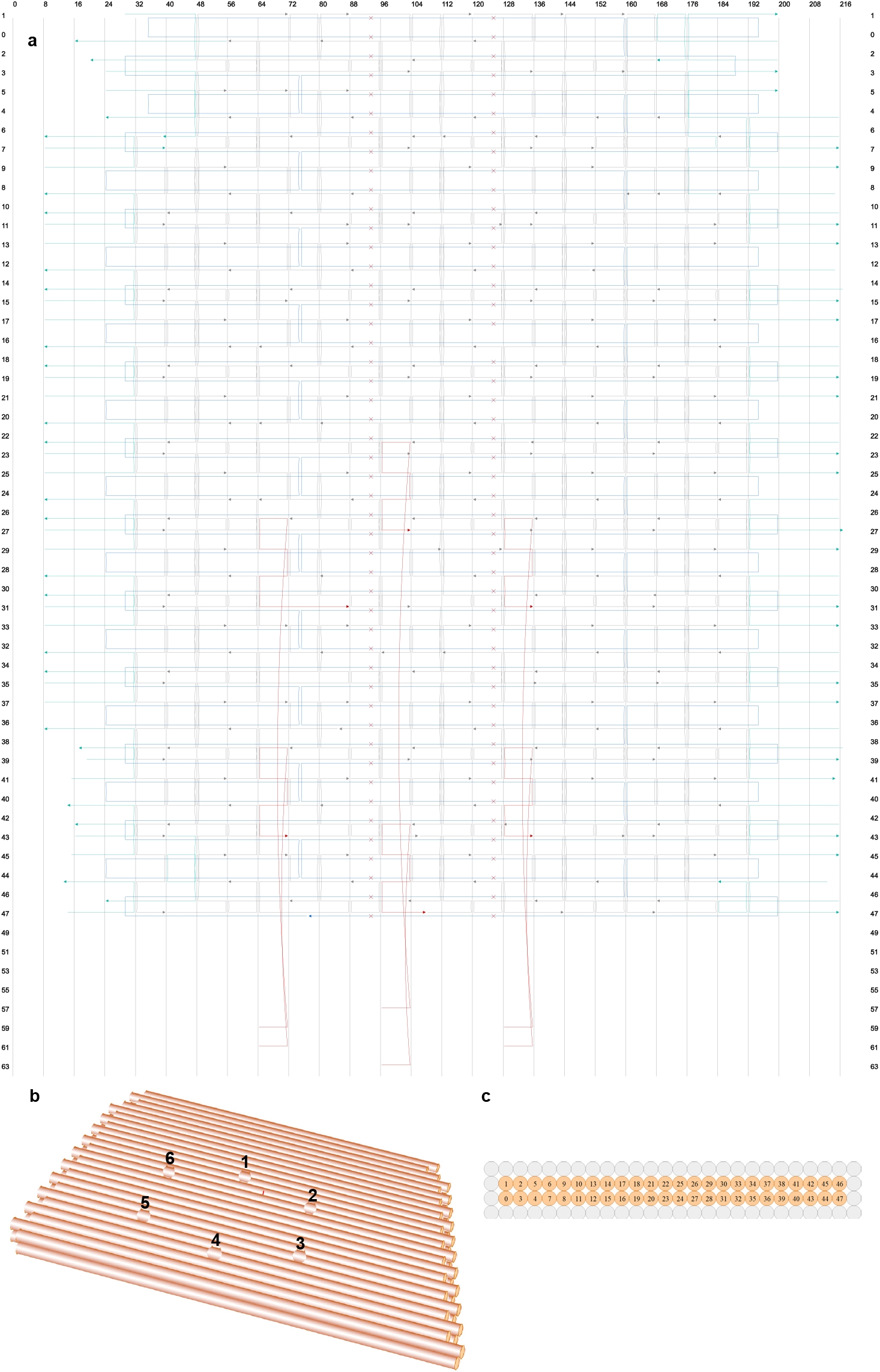
Design of the template for L11. **a,** the design blueprint of the template in cadnanoSQ. The scaffold DNA is in blue, while DNA staples are in gray. The DNA staples colored in cyan have polyA sequences at their non-hybridized parts, which can help reducing aggregation. The 6 staples having protruding sequences at their 5 primes, for the docking of DNA-peptide conjugate, were colored in red. **b,** the 3D view of the tempalte, showing that the 6 protruding sites are presenting a hexagonal pattern, and the distance between adjacent sites are around 10 nm. **c,** the cross-section view of the template in cadnanoSQ.

**Extended Fig.11.**
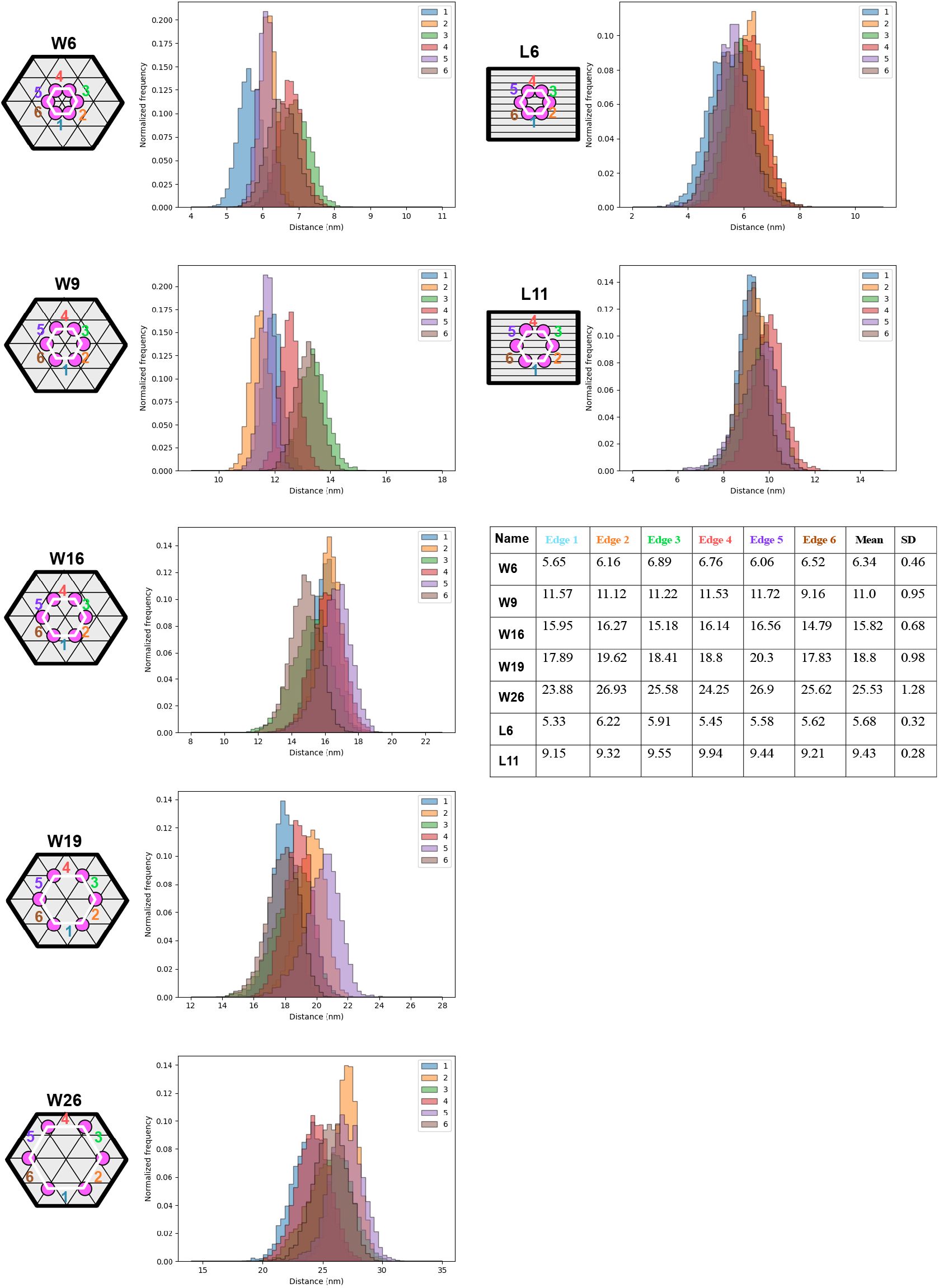
Nucleotide-to-nucleotide distance measurement through oxDNA molecular dynamics simulation. For each hexagonal pattern on DNA origami, the distance distribution (within the oxDNA simulation time frame) of each edge is shown in the distribution histograms. The mean distance of each edge, the mean edge distance of a hexagonal pattern and corresponding standard deviation (SD) are listed in the table.

**Extended Fig.12.**
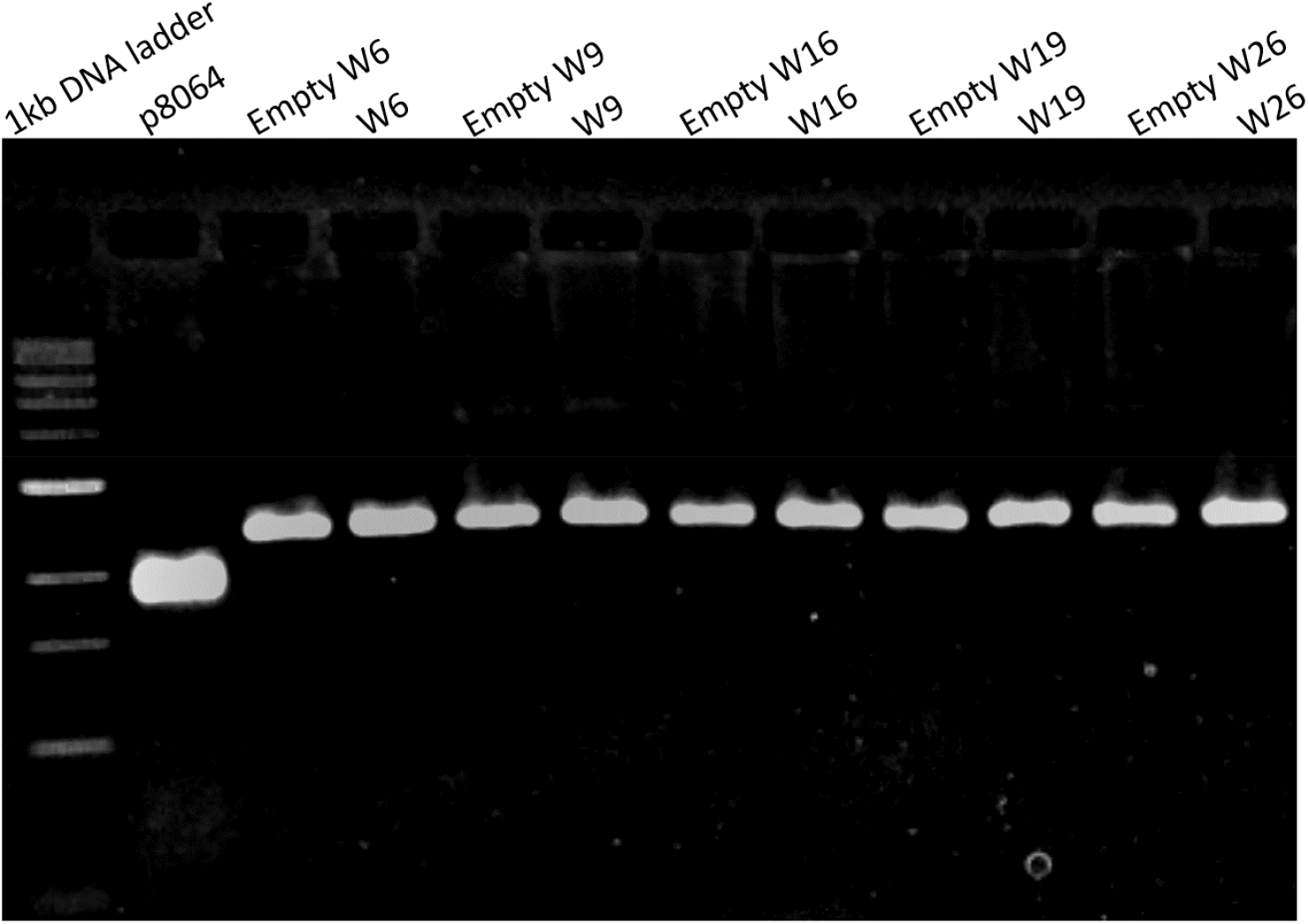
2% agarose gel electrophoresis of all W (with and without peptide attachment) structures. The sample of each well is indicated as in the figure.

**Extended Fig.13.**
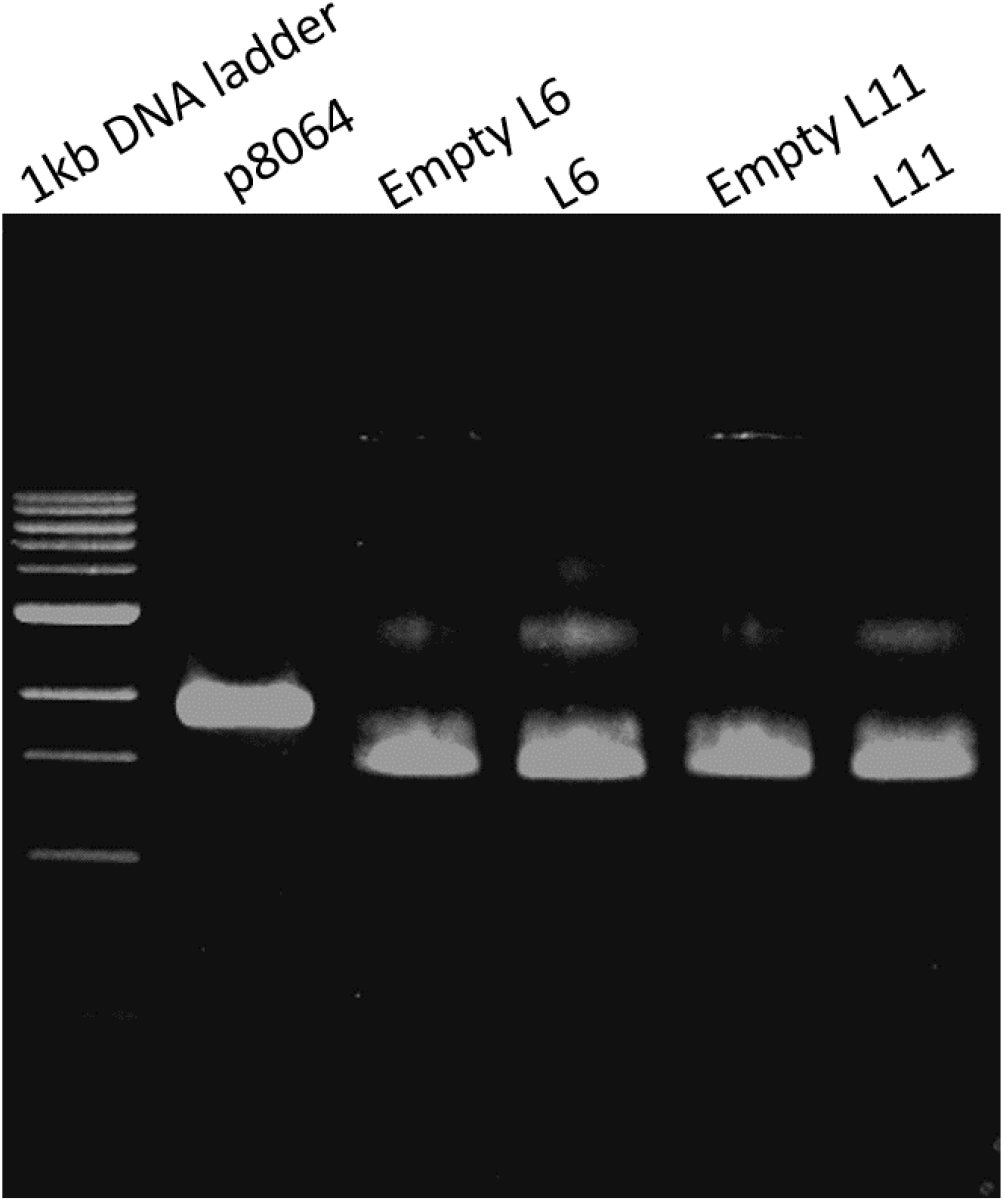
2% agarose gel electrophoresis of all L (with and without peptide attachment) structures. The sample of each well is indicated as in the figure.

**Extended Fig.14.**
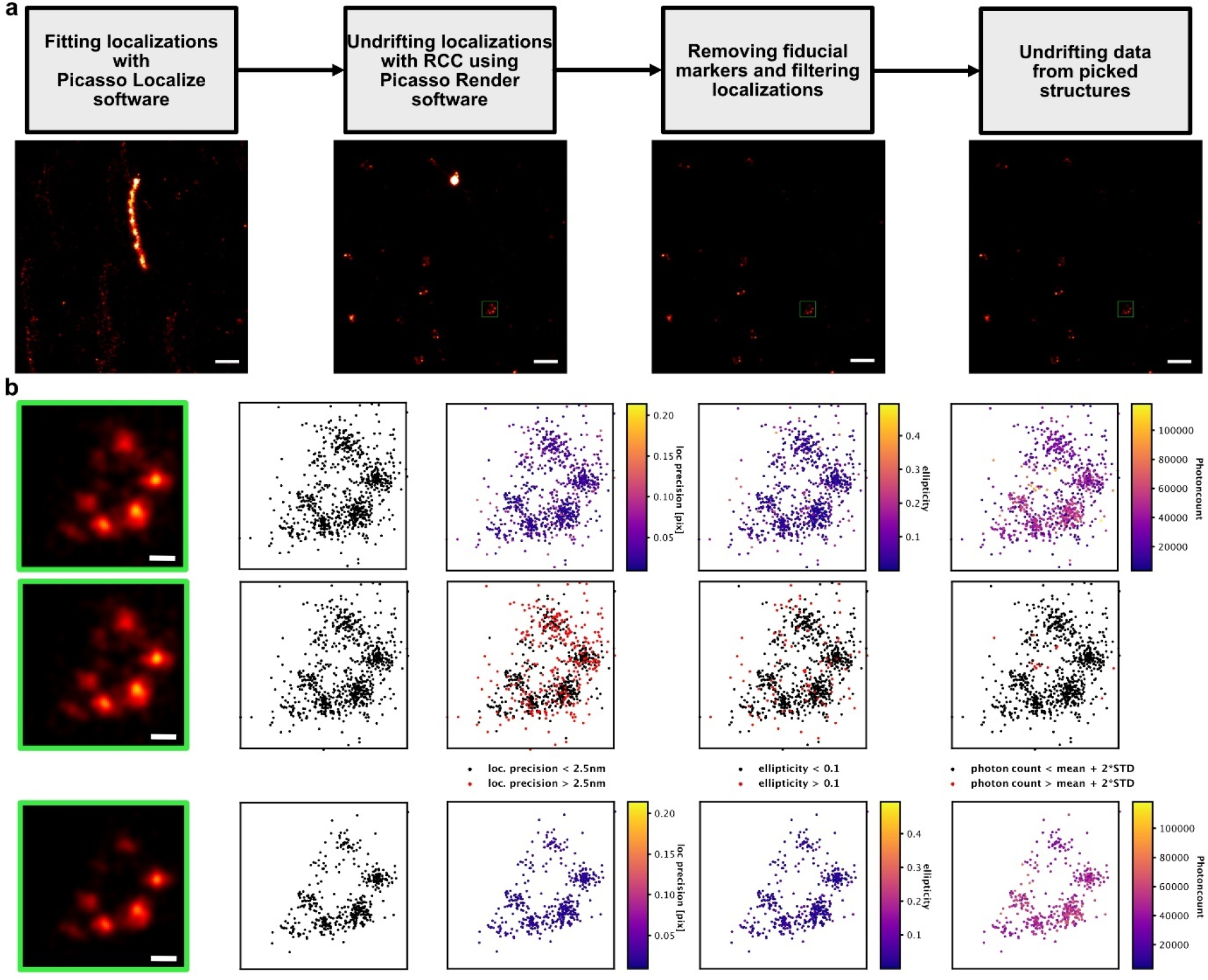
Pre-processing of DNA-PAINT data. **a**, Steps of preprocessing the DNA PAINT data with super resolved field of view images of the resulting DNA PAINT localization data of the TRAIL-peptide functionalized wFS37 structures (scale bar=200nm). **b**, Plots of localizations belonging to a TRAIL-peptide functionalized wFS37 structure taken from the FOV image (green rectangle) showing localization properties before filtering (upper row), localizations removed by the corresponding filters (red) (middle row) and localization properties after filtering (bottom row) along the super resolution image rendered from the corresponding localizations (scale bar = 20nm).

**Extended Fig.15.**
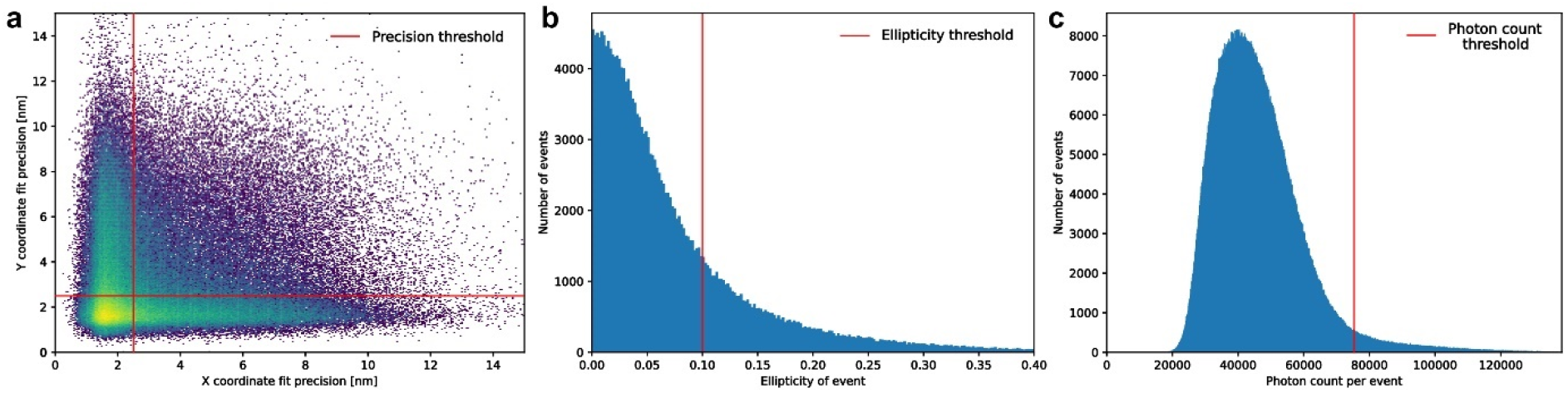
Plots of localization properties and their respective cutoffs used for filtering. **a**, Plots of localization fitting precision, **b**, ellipticity, **c**, and photon counts of the localization data produced by imaging TRAIL-peptide functionalized wFS37 structure with the thresholds applied during preprocessing indicated.

**Extended Fig.16.**
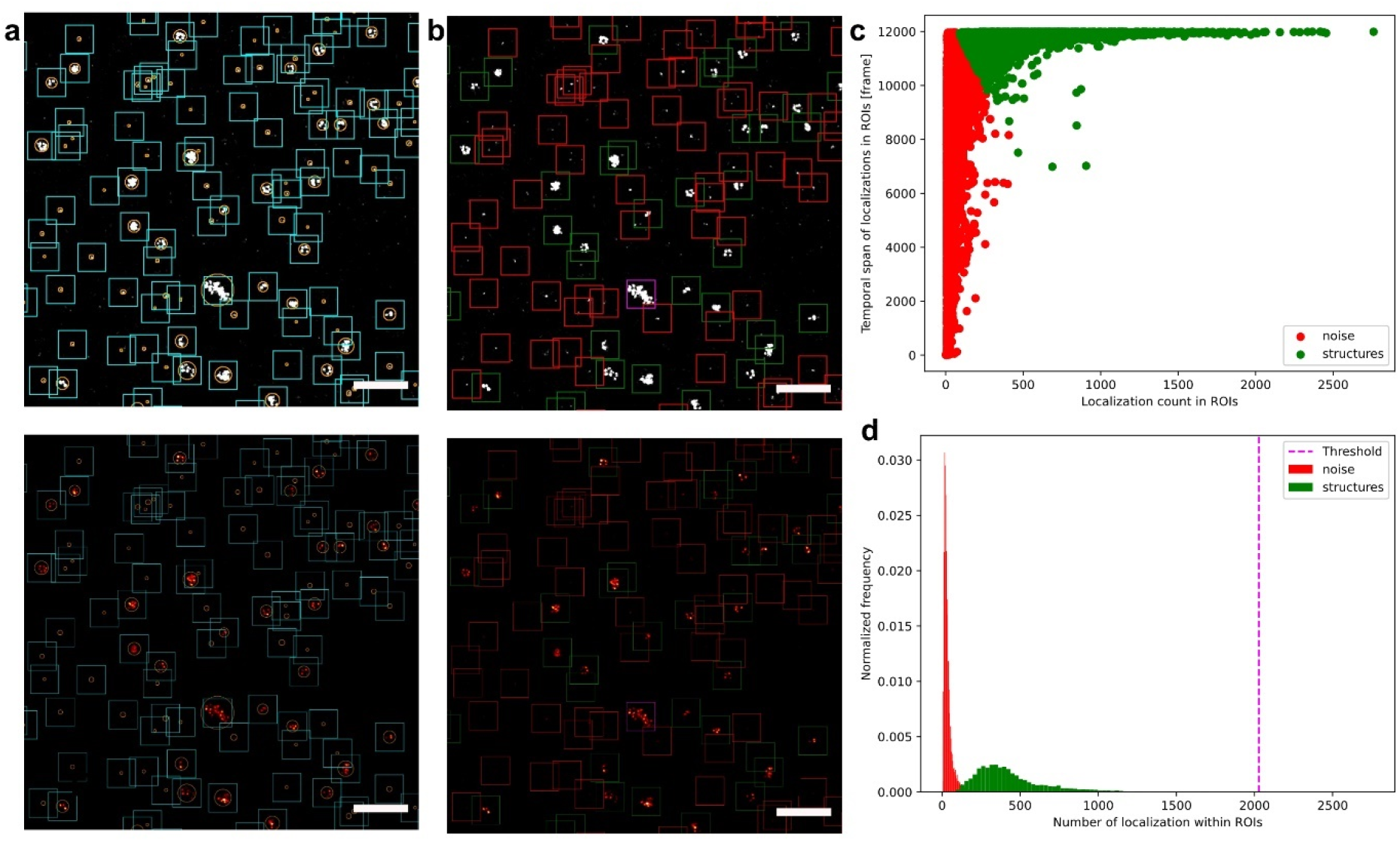
Detection of DNA origami ROI coordinates from DNA-PAINT data (A-D) Steps for DNA origami structure position detection in DNA PAINT data presented on the gray scale (top row) and colormap FOV images (bottom row) of W37 structures (scalebars = 500nm). **a**, Possible positions of origamis structures were found by detecting contours (orange circles) in the grayscale low resolution images rendered from the data and uniform origami ROI coordinates (cyan boxes) were calculated using the centroid coordinates of the contours. **b**, The origami ROIs containing noise (red boxes) were filtered out based on the number and temporal span of localization in them, ROIs containing aggregates of structures (magenta boxes) were filtered out by removing ROIs containing 10 STD more localization than the mean of non-noise containing ROIs, the remaining ROIs (green boxes) were kept as origami ROIs for further processing. **c**, Scatter plot of ROI localization numbers and temporal spans clustered into 2 groups using k-means clustering on two-component PCA transformed data resulting in a cluster of ROIs with shorter temporal span containing noise (red) and ROIs containing signal from structures. **d**, Distribution plot of localization numbers of clustered ROIs showing the localization number threshold (magenta dashed line) above which ROIs containing aggregates of structures were filtered out.

**Extended Fig.17.**
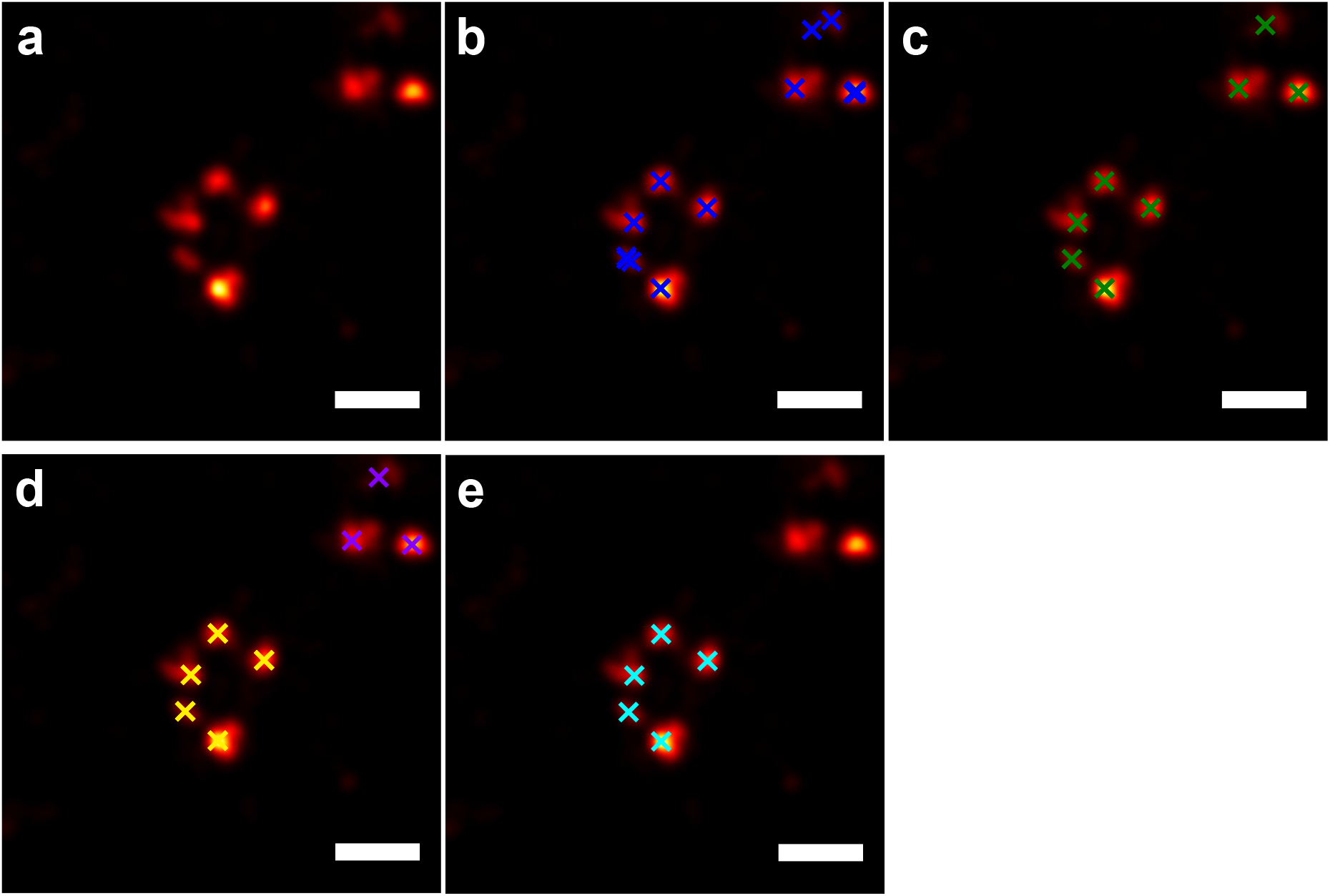
Detection of TRAIL peptides in origami ROIs extracted from DNA-PAINT data. **a**, Localizations lying within detected origami ROIs were grouped and rendered. **b**, Local maxima (blue crosses) were detected in the rendered image. **c**, Maxima lying closer to each other than a distance threshold (0.45 site distance) were merged and an average position for them were calculated yielding the estimated position of TRAIL peptides in the ROI (green crosses). **d**, TRAIL peptides belonging to individual origamis were grouped together (purple and yellow crosses) via hierarchical clustering with the maximum intra cluster distance set to 2.5 site distance. **e**, Finally the cluster with the most number of TRAIL peptides closest to the center of the origami ROI is kept and used for quantification. (Scale bars = 50nm).

**Extended Fig.18.**
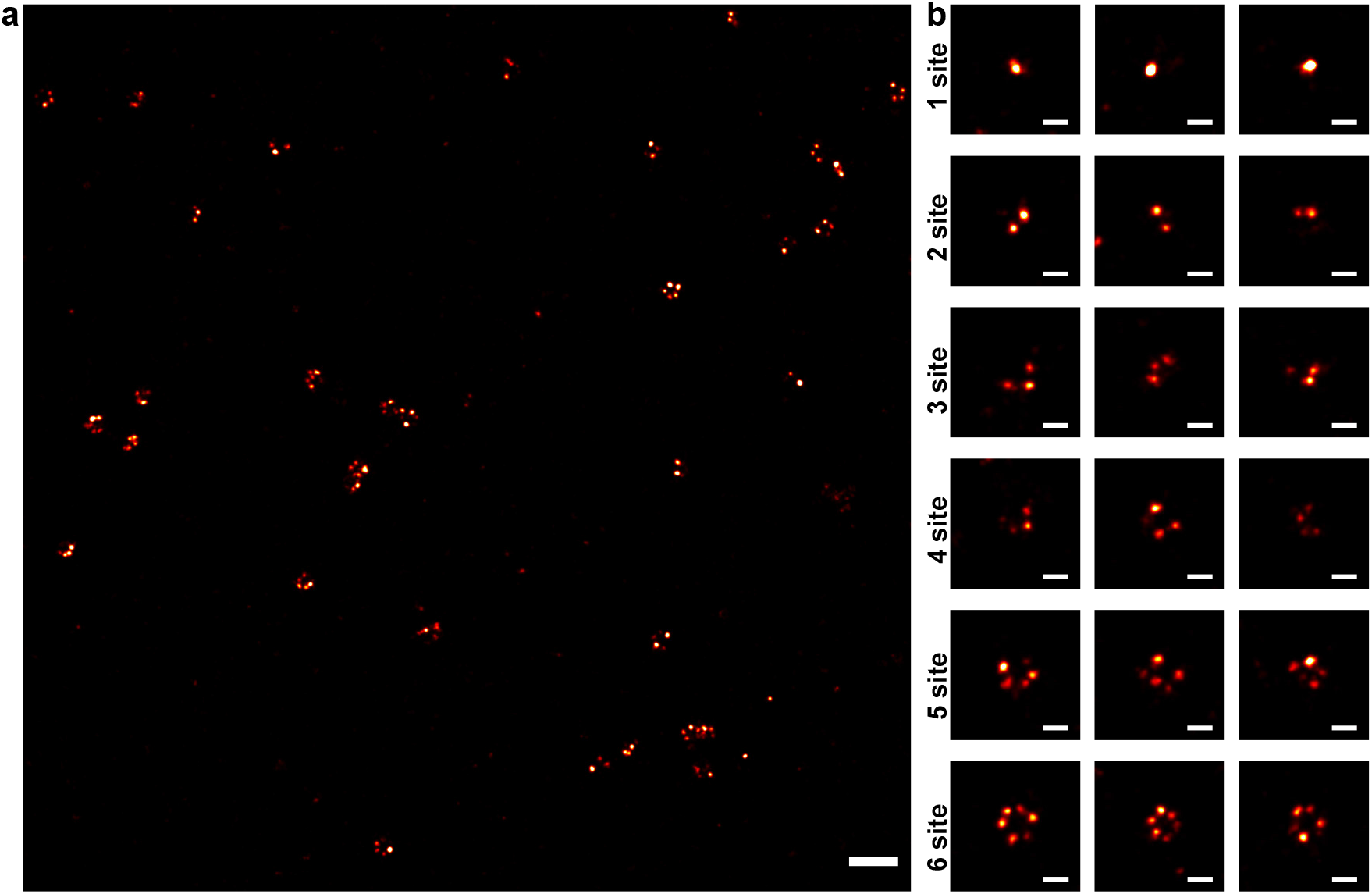
DNA-PAINT images of W37 structures. **a**, A field of view DNA PAINT image of the W37 structures (Scale bar = 200nm). **b**, Rendered images of origami ROIs containing 1, 2, 3, 4, 5 and 6 detected TRAIL peptides (Scale bar = 50nm).

**Extended Fig.19.**
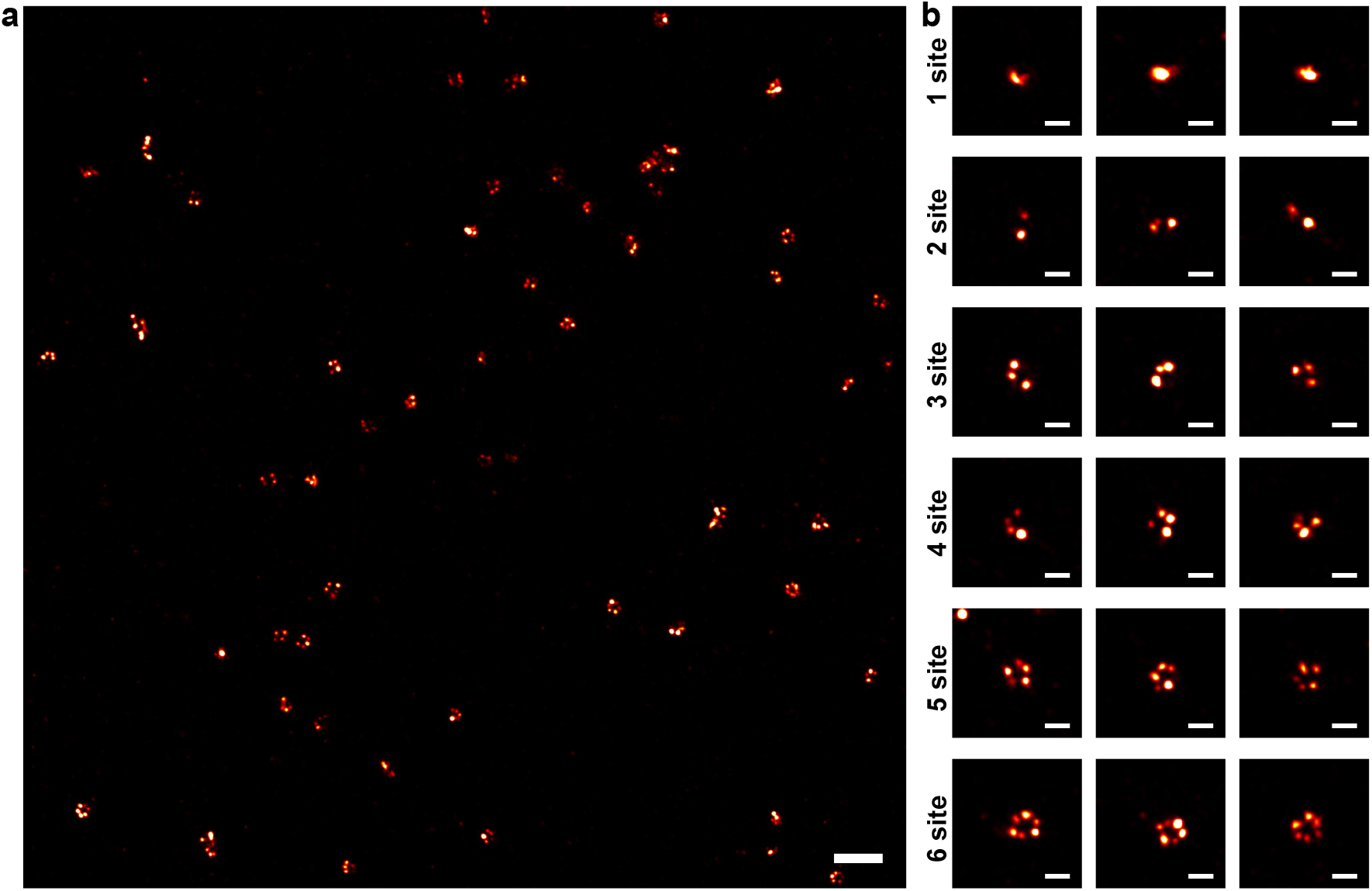
DNA-PAINT images of W28 structures. **a**, A field of view DNA PAINT image of the W28 structures (Scale bar = 200nm). **b**, Rendered images of origami ROIs containing 1, 2, 3, 4, 5 and 6 detected TRAIL peptides (Scale bar = 50nm).

**Extended Fig.20.**
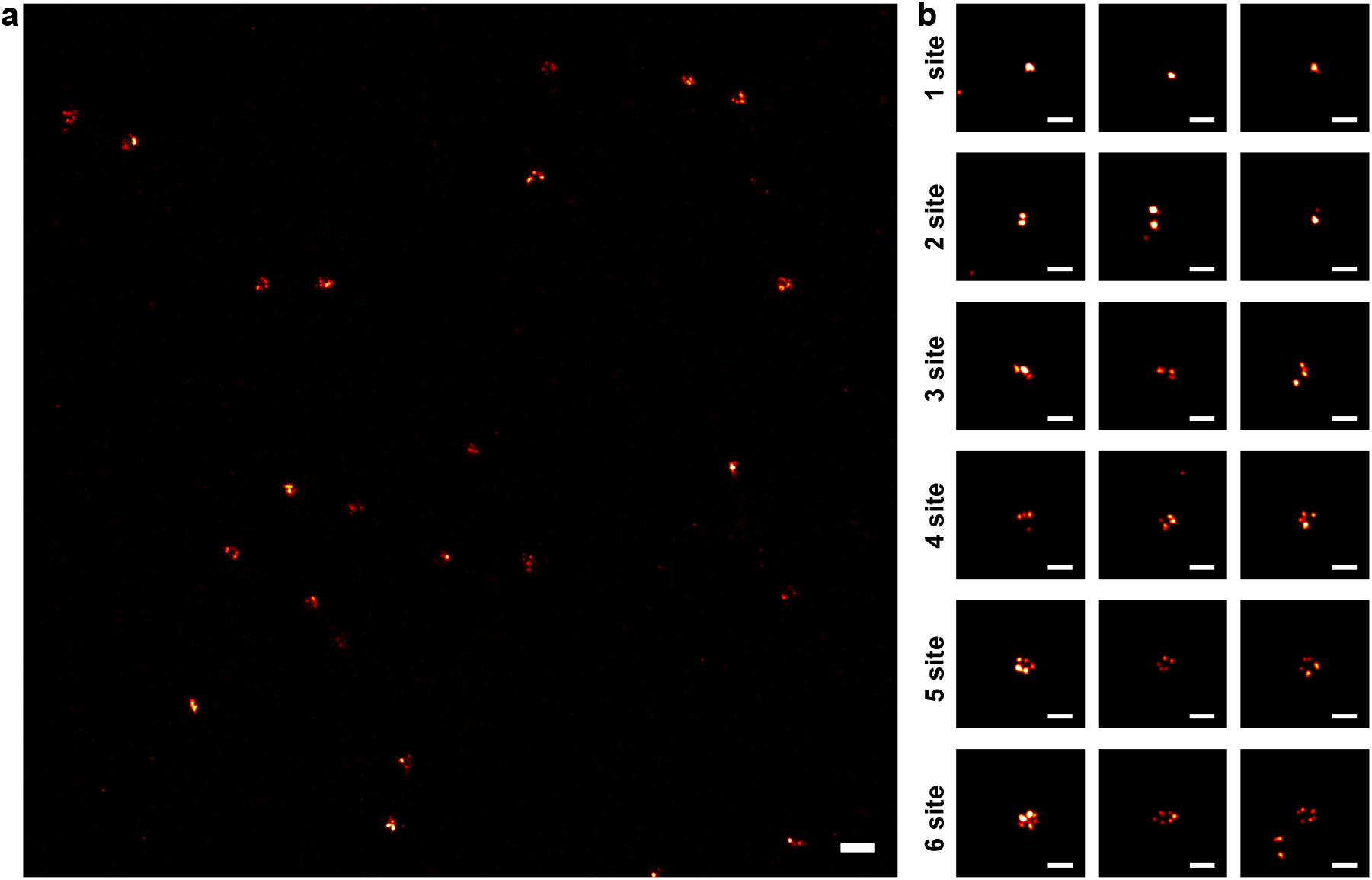
DNA-PAINT images of W19 structures. **a**, A field of view DNA PAINT image of the W19 structures (Scale bar = 200nm). **b**, Rendered images of origami ROIs containing 1, 2, 3, 4, 5 and 6 detected TRAIL peptides (Scale bar = 50nm).

**Extended Fig.21.**
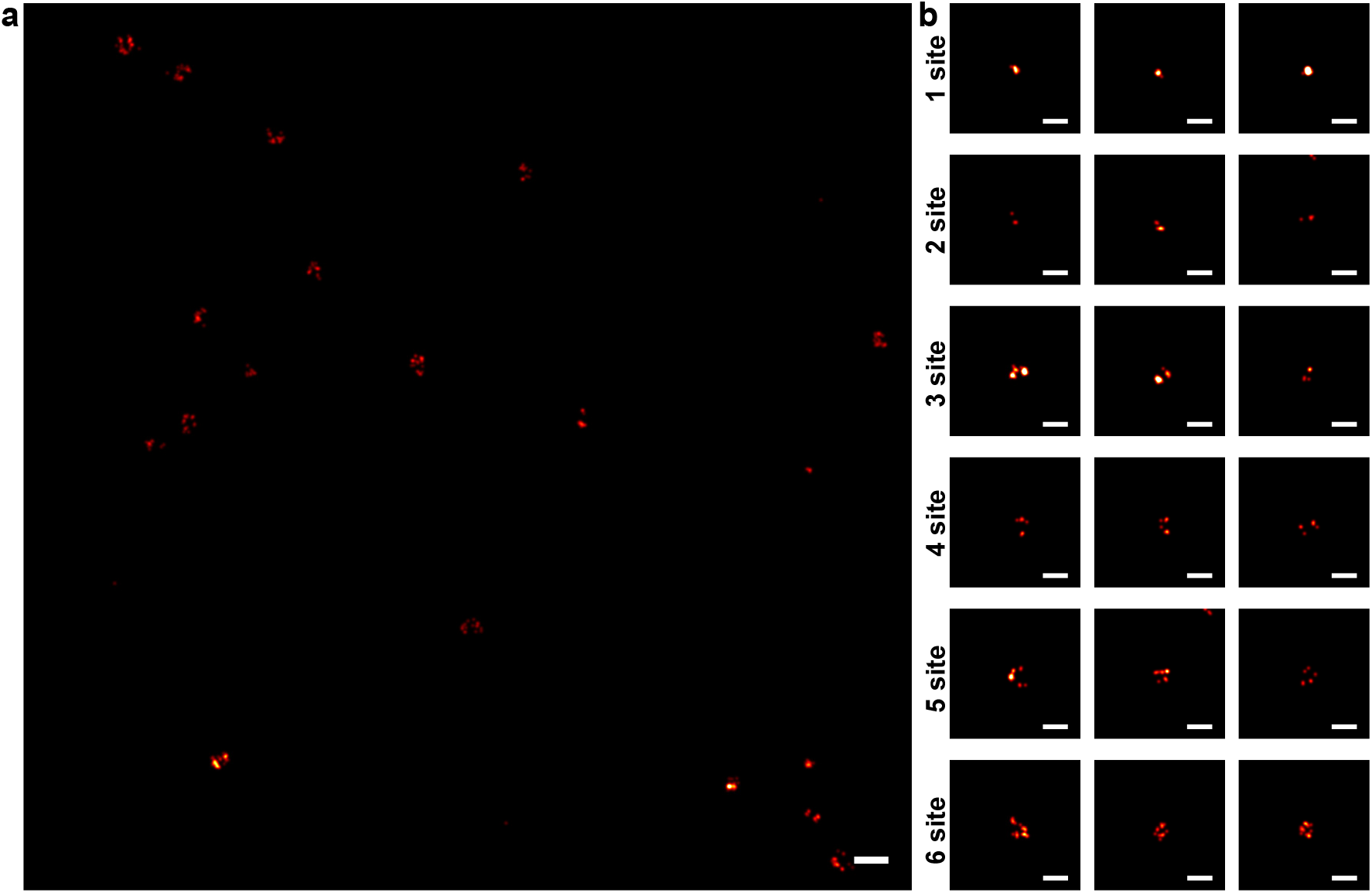
DNA-PAINT images of W16 structures. **a**, A field of view DNA PAINT image of the W16 structures (Scale bar = 200nm). **b**, Rendered images of origami ROIs containing 1, 2, 3, 4, 5 and 6 detected TRAIL peptides (Scale bar = 50nm).

**Extended Fig.22.**
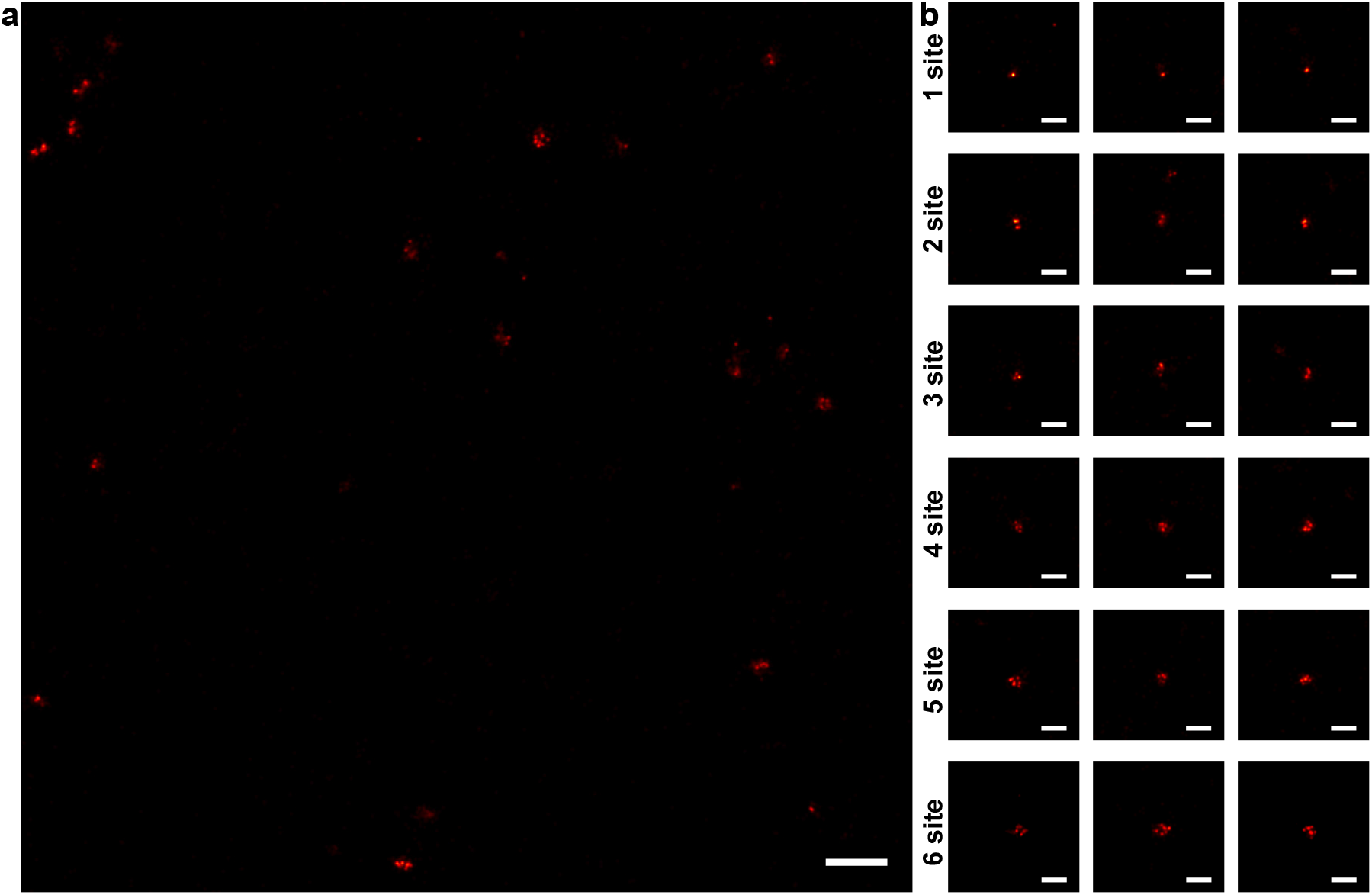
DNA-PAINT images of W9 structures. **a**, A field of view DNA PAINT image of the W9 structures (Scale bar = 200nm). **b**, Rendered images of origami ROIs containing 1, 2, 3, 4, 5 and 6 detected TRAIL peptides (Scale bar = 50nm).

**Extended Fig.23.**
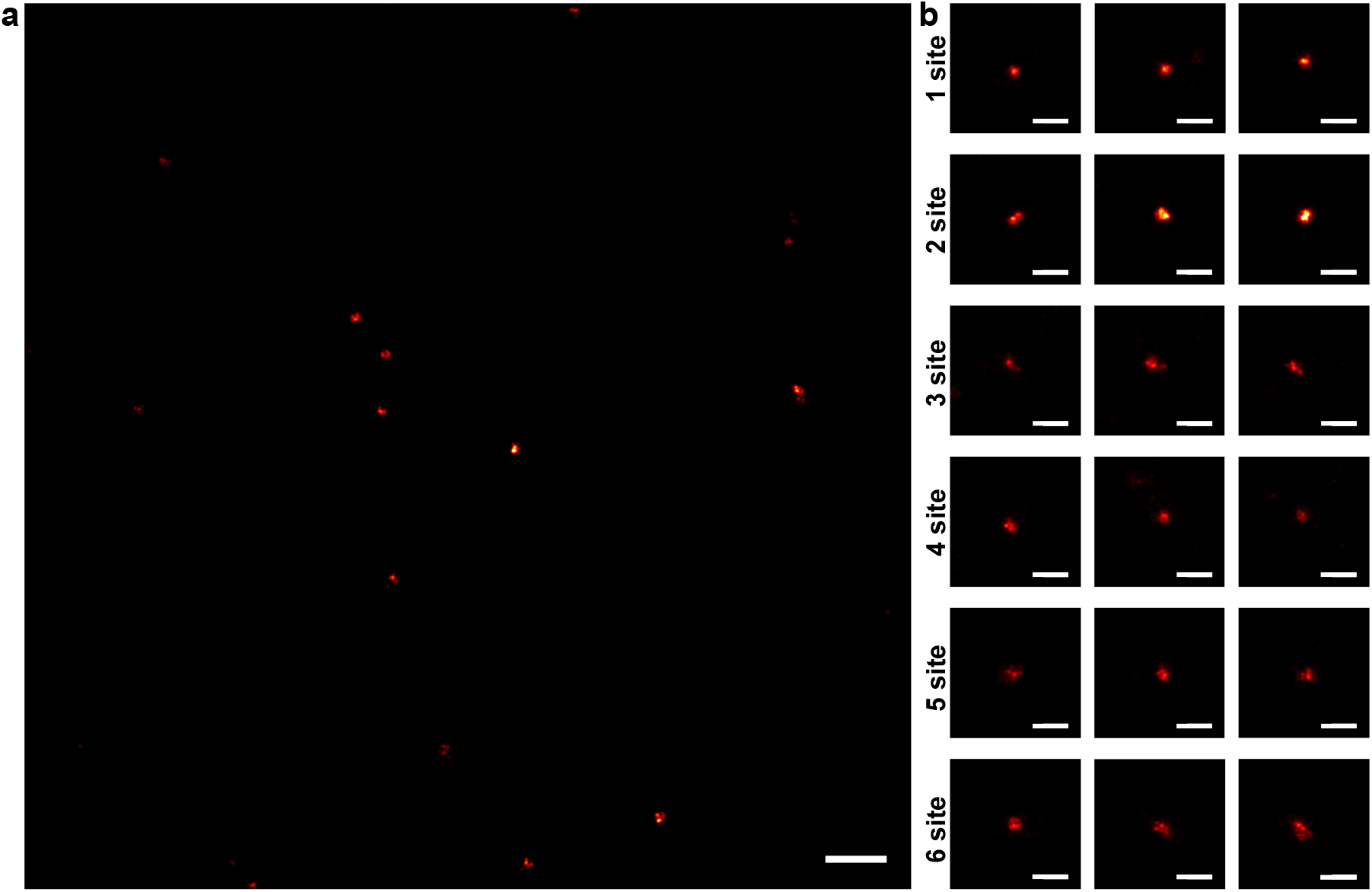
DNA-PAINT images of W6 structures. **a**, A field of view DNA PAINT image of the W6 structures (Scale bar = 200nm). **b**, Rendered images of origami ROIs containing 1, 2, 3, 4, 5 and 6 detected TRAIL peptides (Scale bar = 50nm).

**Extended Fig.24.**
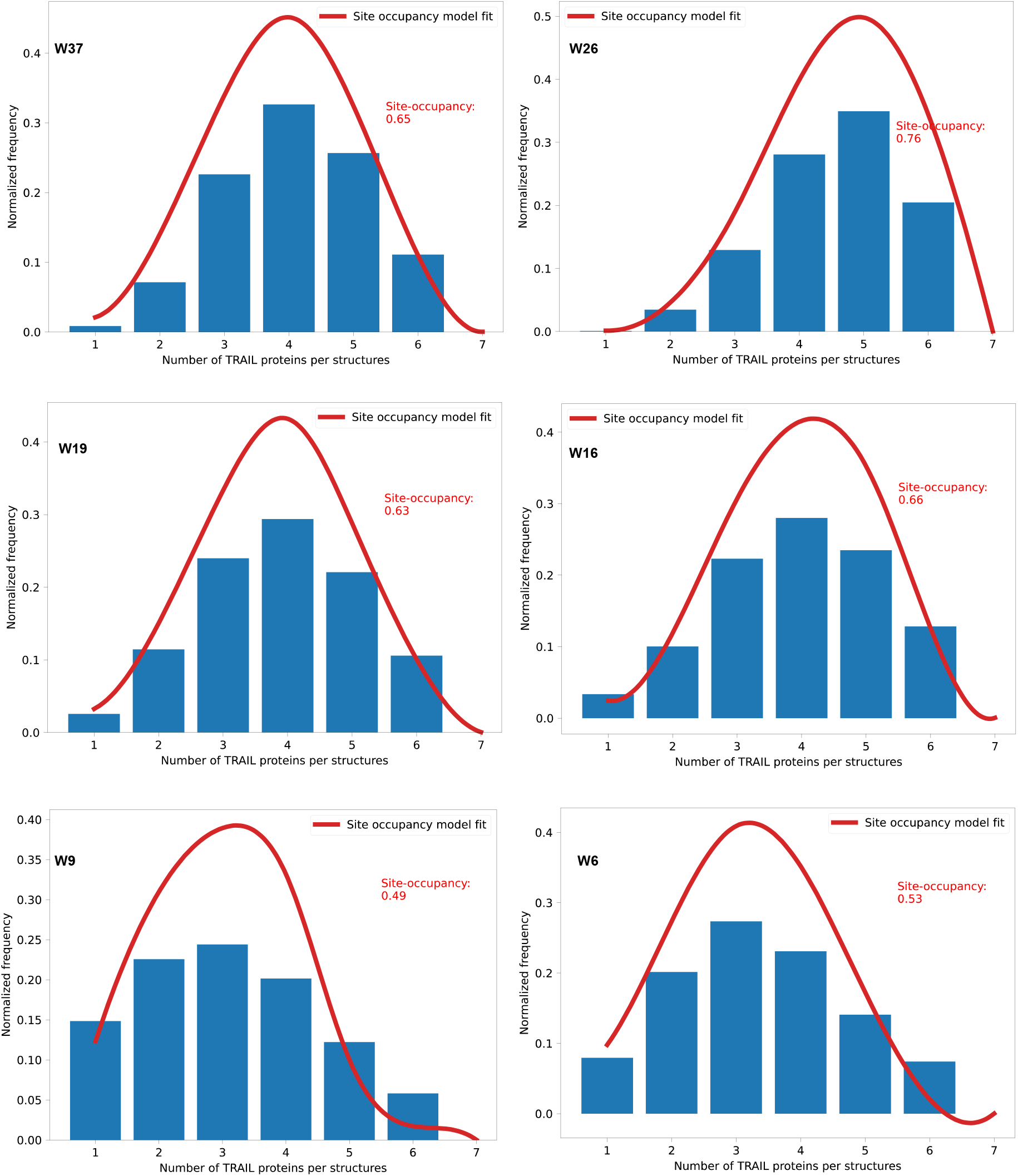
Protruding DNA site occupancy and the distribution of number of peptide per structure for each W-structure. Computed from the DNA-PAINT data.

**Extended Fig.25.**
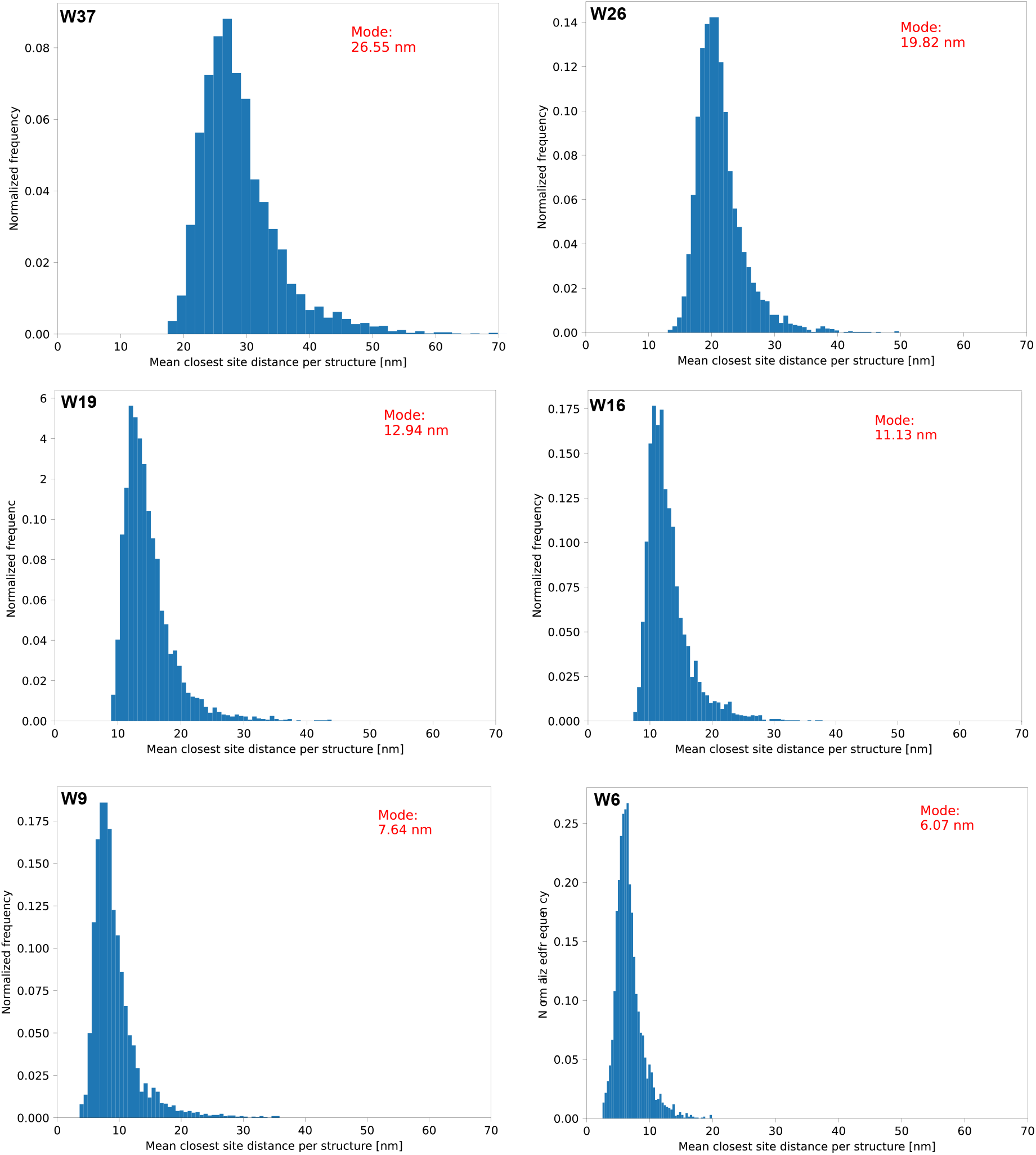
The inter-peptide distance distribution of each W-structure. Computed from the DNA-PAINT data.

**Extended Fig.26.**
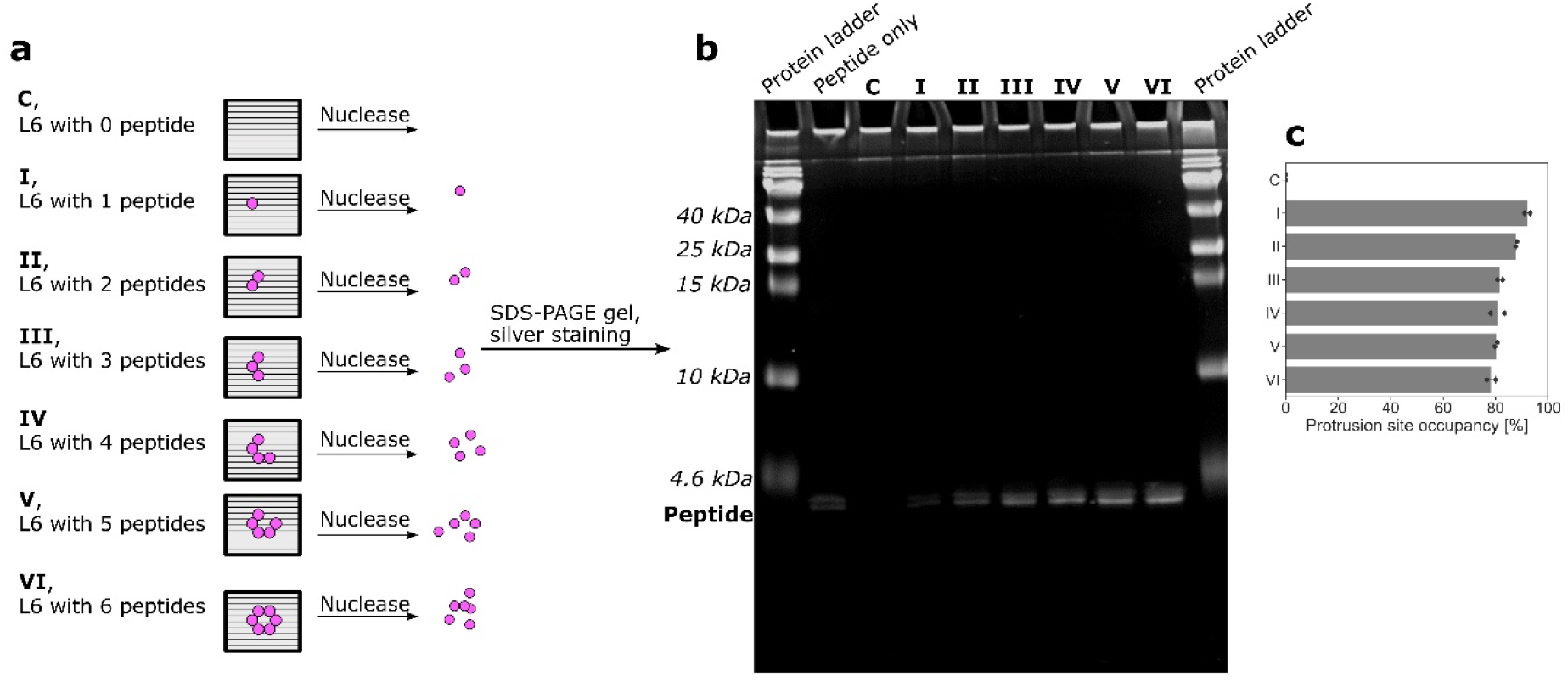
Gel-based peptide quantification of L6. L6 structures attached with different numbers of peptides (as indicated in **a**) were prepared, then they were treated with DNase I to completely digest the DNA origami template. The samples were then run on SDS-PAGE (4-20% gradient gel) gel. Silver staining was carried out to visualize the peptide (as indicated in **b**, the bands below 4.6 kDa ladder)). We did not see obvious DNase I band, which could be caused by its very low loading amount. The protrusion site occupancy calculated from the band intensities plotted in **c**.

**Extended Fig.27.**
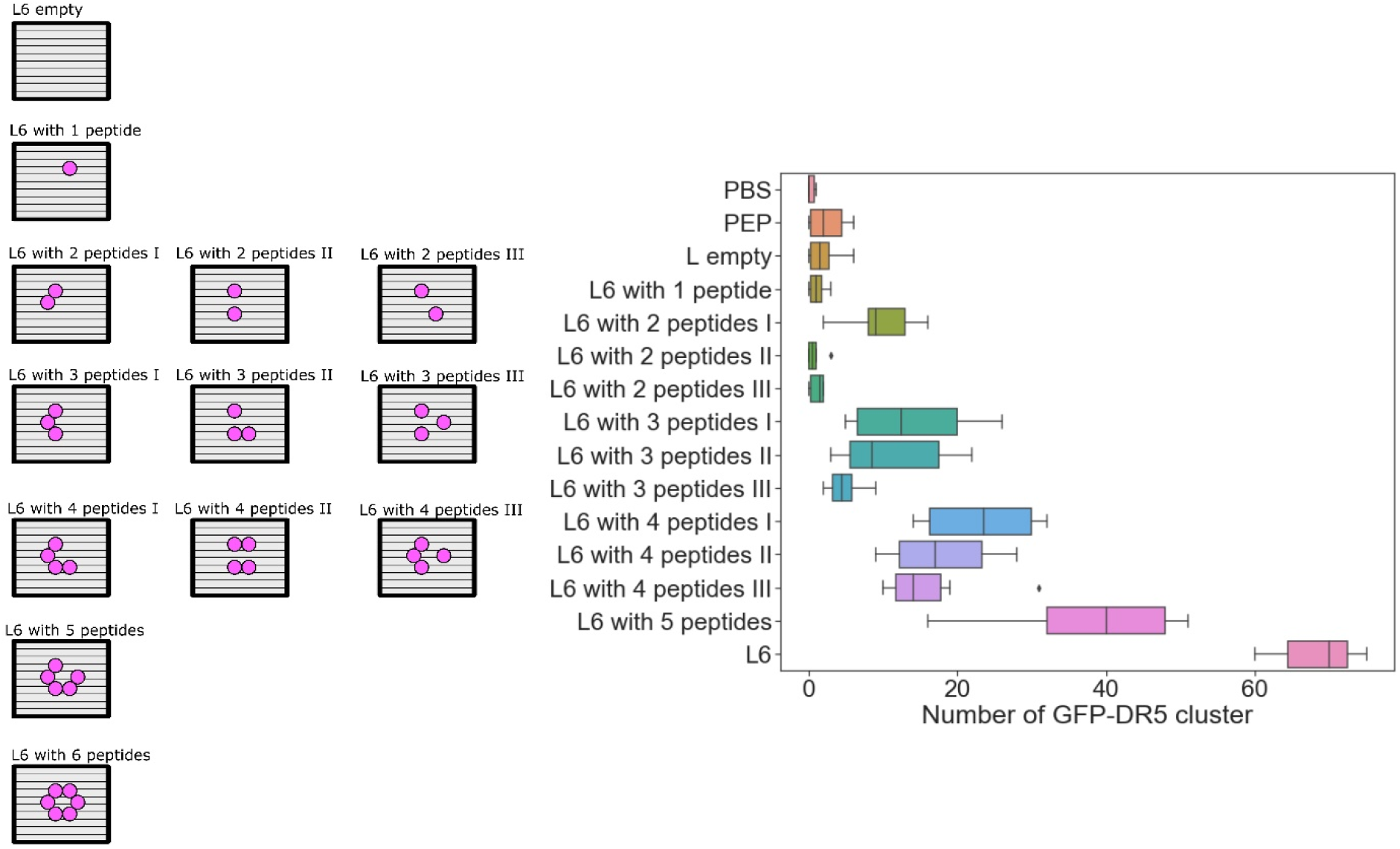
Quantification of GFP-DR5 clusters induced by L6 attached with different number of peptides on GFP-DR5-expressing MCF-7 cell. Left: the illustration of peptide numbers and patterns; Right: GFP-DR5 cluster counting. (n = 50 cells). The treatment was 4 hours with 2-nM DNA origami structures (12-nM peptides).

**Extended Table.1.** The half maximal inhibitory concentration (IC_50_) of peptides.

| Treatment | $IC_{50}$ [nM] MCF-7 | $IC_{50}$ [nM] SK-BR-3 | $IC_{50}$ [nM] MDA-MB-231 |
| --- | --- | --- | --- |
| **Peptide** | [a] | [a] | [a] |
| **W** | [b] | [b] | [b] |
| **W6** | 14.3 | 9.0 | 9.4 |
| **W9** | 52.7 | 44.2 | 68.3 |
| **W16** | [a] | [a] | [a] |
| **W19** | [a] | [a] | [a] |
| **W26** | [a] | [a] | [a] |
| **L** | [b] | [b] | [b] |
| **L6** | 2.2 | 4.5 | 3.7 |
| **L11** | [b] | [b] | 292.7 |

[a] Determined up to 300 nM of peptide.
[b] Determined up to 50 nM of DNA origami structures.

## Notes

### Competing Interest Statement

The authors have declared no competing interest.

